# Regulation of Src tumor activity by its N-terminal intrinsically disordered region

**DOI:** 10.1101/2021.05.10.443360

**Authors:** Emilie Aponte, Marie Lafitte, Audrey Sirvent, Valérie Simon, Maud Barbery, Elise Fourgous, Mariano Maffei, Florence Armand, Romain Hamelin, Julie Pannequin, Philippe Fort, Miquel Pons, Serge Roche

**Affiliations:** CRBM, CNRS, Univ. Montpellier, F-34000, Montpellier France; Biomolecular NMR laboratory, Department of Inorganic and Organic Chemistry, University of Barcelona, Baldiri Reixac 10-12, 08028, Barcelona, Spain; Proteomics Core Facility, School of Life Sciences, École Polytechnique Fédérale de Lausanne (EPFL), 1015 Lausanne, Switzerland; IGF, CNRS, Univ. Montpellier, F-34000 Montpellier, France; Equipe labellisée Ligue Contre le Cancer, CRBM, CNRS, Univ. Montpellier, F-34000, Montpellier France; Evvivax srl - Via di Castel Romano, 100 - 00128 - Rome, Italy

**Keywords:** cell signaling, intrinsically disordered region, protein kinase, cell transformation

## Abstract

The membrane anchored Src tyrosine kinase is involved in numerous pathways and its deregulation is involved in human cancer. Our knowledge on Src regulation relies on crystallography, which revealed intramolecular interactions to control active Src conformations. However, Src contains a N-terminal intrinsically disordered unique domain (UD) whose function remains unclear. Using NMR, we reported that UD forms an intramolecular fuzzy complex involving a conserved region with lipid-binding capacity named Unique Lipid Binding Region (ULBR), which could modulate Src membrane anchoring. Here we show that the ULBR is essential for Src’s oncogenic capacity. ULBR inactive mutations inhibited Src transforming activity in NIH3T3 cells and in human colon cancer cells. It also reduced Src-induced tumor development in nude mice. An intact ULBR was required for MAPK signaling without affecting Src kinase activity nor sub-cellular localization. Phospho-proteomic analyses revealed that, while not impacting on the global tyrosine phospho-proteome in colon cancer cells, this region modulates phosphorylation of specific membrane-localized tyrosine kinases needed for Src oncogenic signaling, including EPHA2 and Fyn. Collectively, this study reveals an important role of this intrinsically disordered region in malignant cell transformation and suggests a novel layer of Src regulation by this unique region via membrane substrate phosphorylation.

## INTRODUCTION

Src, originally identified as an oncogene, is a membrane-anchored tyrosine kinase, which mediates signaling induced by a wide range of cell surface receptors, leading to cell growth and adhesion (1). Src deregulation is associated with cancer development, although the underlying mechanisms are not fully understood (2,3). Src shares with the other Src Family Kinases (SFKs) a common modular structure formed by the membrane-anchoring SH4 region followed by an intrinsically disordered region (IDR) named unique domain (UD), and the SH3, SH2, and kinase domains (2). Our knowledge of Src regulation relies on crystallographic data that revealed SH2 and SH3-dependent intramolecular interactions that control Src catalytic activity (4). However, the functions of the SH4-UD module have often been disregarded because of X-ray invisibility. UD is the mostly divergent part of SFK proteins, which supported the idea of a unique function among SFKs (5,6). However, early studies reported that the whole UD deletion does not affect Src oncogenic activity (7), which suggests that this region may not play an important role in Src signaling. While considerable insight into Src regulation has been provided since the discovery of the Src oncogene, the functional role of its unstructured region remains unclear.

IDRs are highly prevalent in proteins regulating essential cell processes, such as transcription or signaling that are implicated in human diseases (8). The integration of multiple weak interactions is crucial for the increasingly recognized role of IDRs in the formation of membrane-less organelles, through liquid-liquid phase separation (8). Multiple, rapidly exchanging weak contacts are also at the basis of the formation of the so called “fuzzy complexes” by IDRs, in which the IDR remains disordered, but the complex is stabilized by multiple transient contacts (5,9). Intramolecular fuzzy complexes involve a similar fuzzy interaction between an IDR and a globular domain in the same protein to form a relatively compact structure (5,9). Intramolecular fuzzy complexes may regulate the communication between the disordered and globular regions of a signaling protein and sense the environment (e.g. membrane lipids) that will influence the activity of folded domains, such as kinase activity or the binding capacity of SH2 and SH3 domains (5,9). Thus, IDRs provide a unique mechanism of protein regulation by the local environment. In line with this, recent molecular studies uncovered key features of such a new Src-UD regulatory mechanism (10–13). Specifically, Src-UD forms an intramolecular fuzzy complex, where its conformational freedom is restricted by multiple contacts with the globular SH3 domain (11). Although their primary sequences are divergent, UD fuzzy complexes are present in other SFKs, suggesting that this IDR defines a central regulatory mechanism in SFKs (14,15). Our molecular studies on Src identified a conserved region that contributes to the interaction of the UD with the RT and nSrc loops of the SH3 domain (10,11). This region displays affinity with phospholipids and was named Unique Lipid Binding Region (ULBR)(10). Further *in vitro* studies showed that ULBR participates in the interaction between the N-terminal myristoyl group and the SH3 domain and contributes to the modulation of membrane anchoring (13), suggesting that it could also be involved in substrate selection and signaling. Here we start addressing the oncogenic relevance of the Src UD by focusing on ULBR.

## RESULTS

### Evolutionary conservation of ULBR in SFK-UD

Although Src-UD is thought to be evolutionary conserved, we noticed a strong sequence divergence between mammalian and invertebrates SFKs. To clarify this point, we readdressed SFK sequence conservation by comparing the selective constraints exerted along SFK sequences. We computed ω, the ratio of non-synonymous (dN) to silent (dS) mutations from a multiple alignment of Src, Yes and Fyn coding sequences of seven primate species, covering 74 million years of evolution. In such framework, neutral selection is associated with an ω-value close to 1, while values <1 or >1 indicate purifying and diversifying selection, respectively. As shown in Figure 1a, all domains produced ω-values <1. However, UD ω-values indicated variable selection levels depending on SFK members. Notably, the heaviest selection was exerted on Fyn UD (ω = 0.0001), while the selection exerted on Src UD was moderate (ω = 0.126) and even lighter for Yes (ω = 0.363). This indicates that UD sequences are not under neutral selection and supports the notion that these sequences play essential roles in SFKs functions. Consistent with this idea, a comparative sequence analysis of SFK-UD IDR sequences from 10 vertebrate species revealed that residues 64-Phe-Gly-Gly-66 (human Src numbering) are highly conserved across species and also across the three functional related Src, Fyn and Yes. They are part of the conserved Src ULBR (residues 60-67) highlighting an important function for this small region (Fig 1b).

**Figure 1.**
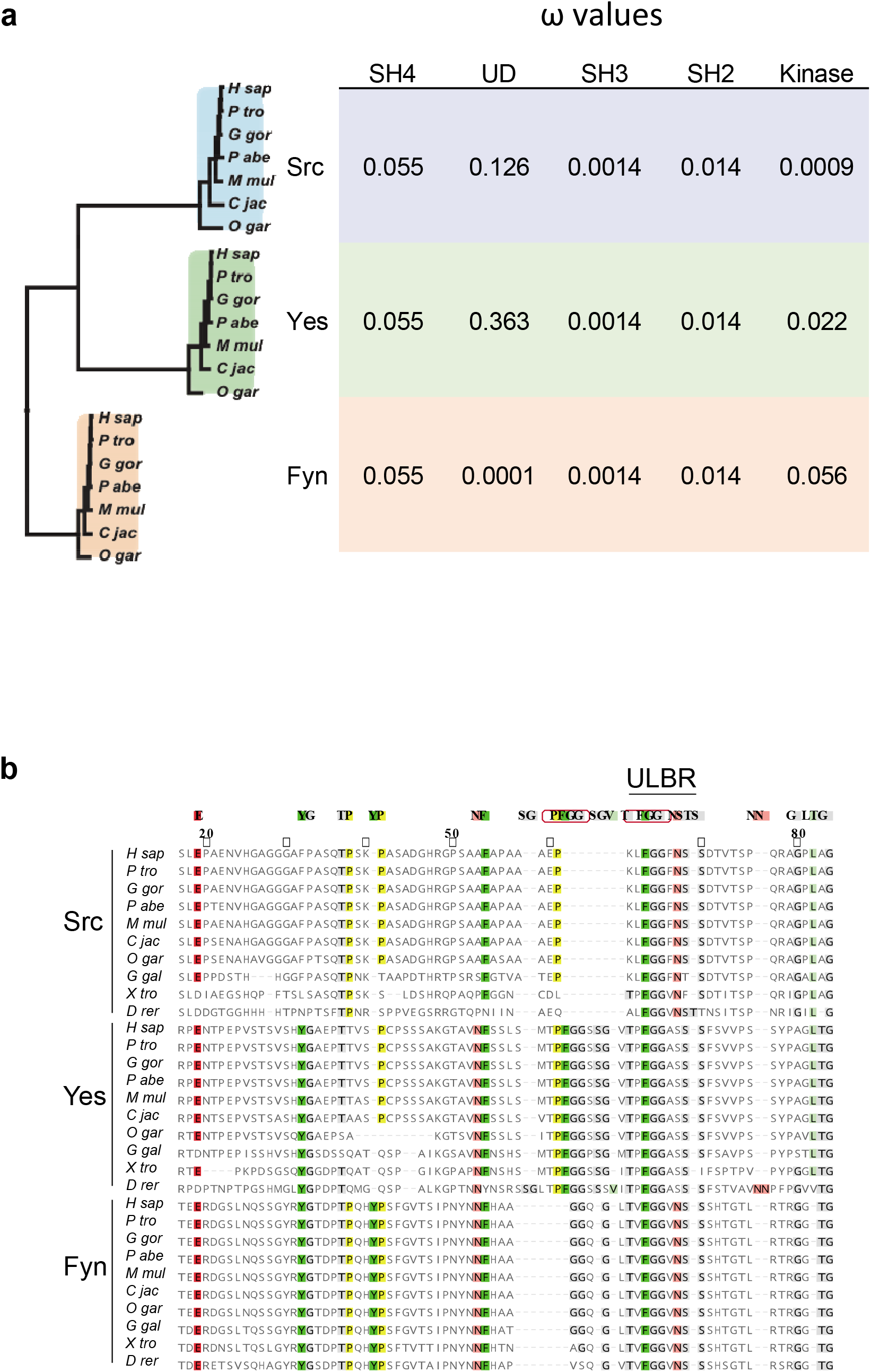
Evolutionary conservation of the UD in SFKs. **a:** Analysis of dN/dS ratios (ω) of the SH4, UD, SH3, SH2 and kinase domains in primate SFKs. Nucleic sequences coding the different domains were aligned based on translation and processed by codeml to estimated ω-values. The phylogenetic tree on the left was generated by PhyML analysis of a multiple alignment of full-length SFK sequences. **b**: Multiple alignment of UD sequences of primate SFKs. Sequences were aligned with MAFFT. The most conserved residues are indicated above the alignment. The strictly conserved FGG of the ULBR is framed. Numbering corresponds to human c-Src sequence. H sap: Homo sapiens; P tro: *Pan troglodytes*; G gor: *Gorilla gorilla*; P abe: *Pongo abelii*; M mul: *Macaca mulatta*; C jac: *Callithrix jacchus*; O gar: *Otolemur garnettii*; G gal: *Gallus gallus*; X tro: *Xenopus tropicalis*; D rer: *Danio rerio*.

### ULBR inactivation affects Src oncogenic activity

We next explored the functional role of Src-ULBR on cell transformation. For this, ULBR was inactivated by replacement of residues 63–65 (Leu-Phe-Gly) by three alanines (named Src3A), known to affect Src-UD molecular properties (i.e. binding to phospholipids and SH3, Src membrane anchoring) (10,11,13). Since these ULBR properties can be regulated by phosphorylation of surrounding Ser69 and Ser75 (10,16), we also inactivated ULBR by replacement of Ser69 or Ser75 by the phospho-mimicking glutamic acid (ie SrcS69E and SrcS75E) (Fig 2a). These ULBR mutations were incorporated in the oncogenic SrcY530F mutant, in which the pTyr530-SH2 interaction is destabilized, thereby inducing an active and open Src conformation (4). Transforming activity was then assessed upon retroviral transduction in immortalized mouse embryonic fibroblasts NIH3T3 (Fig 2b & c). SrcY530F protein levels were reduced compared to regulated Src (Fig 2b), which was previously attributed to an autoregulatory mechanism mediated by the substrate and E3 ligase Cbl (17). Despite this and unlike wild-type Src, oncogenic SrcY530F expression induced anchorage-independent growth as measured by the number of colonies in soft agar. Interestingly, ULBR inactive mutations strongly reduced this transforming effect, suggesting that an intact ULBR is required for oncogenic Src activity (Fig 2c). The invasive properties of NIH3T3 cells were also reduced, as assayed in Boyden chambers coated with matrigel (Fig 2d). However, other transformation related properties induced by SrcY530F, such as dissolution of F-actin bundles causing actin cytoskeletal rearrangement (Fig S1c) (18), were retained in ULBR mutants. This indicates that ULBR selectively regulates some of Src transforming activities.

**Figure 2.**
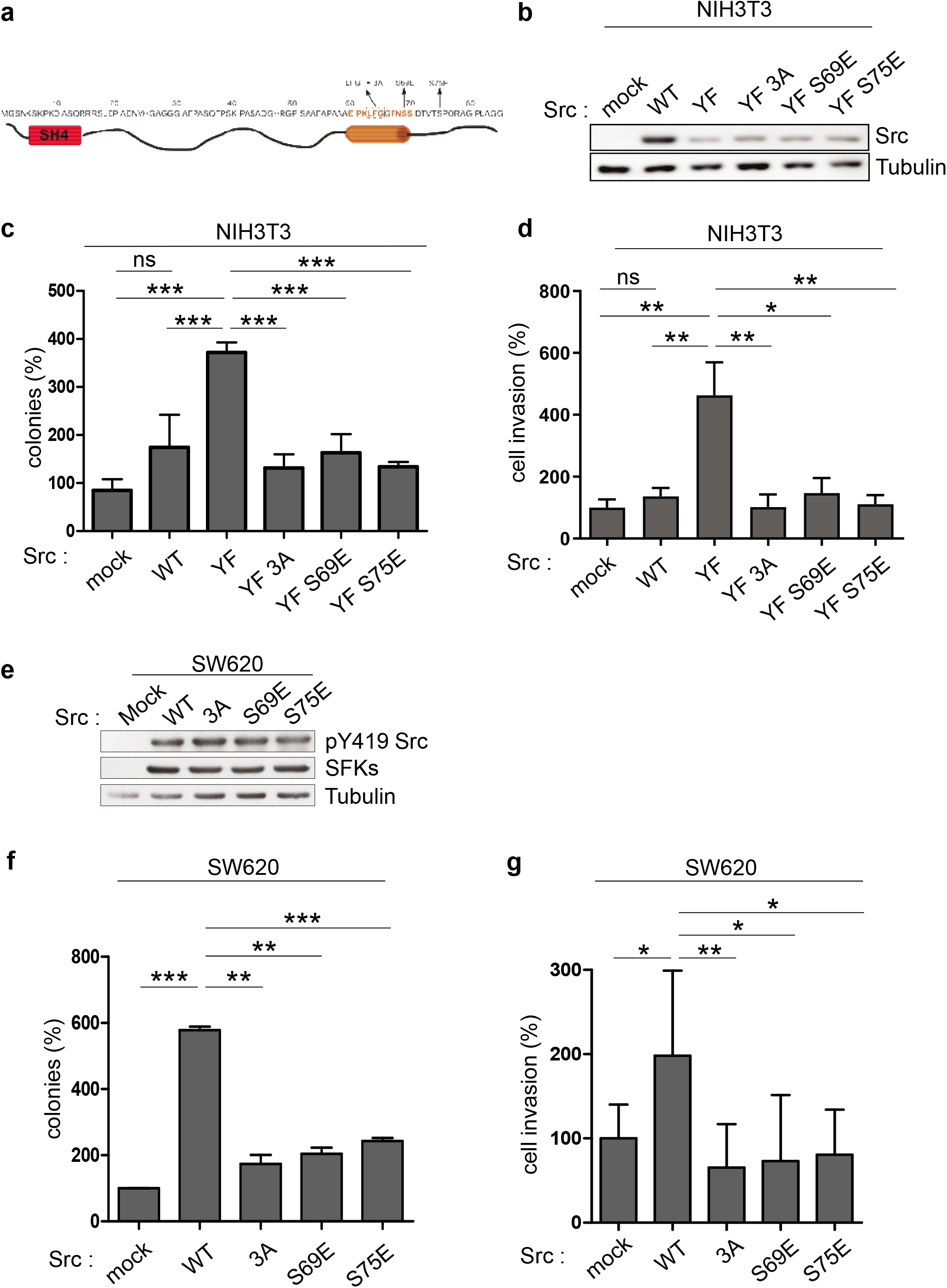
ULBR inactivation affects Src oncogenic activity. **a**: Strategy of ULBR inactivation. **b**: the level of Src expression in NIH3T3 cells transduced with indicated Src constructs. **c, d**: anchorage independent cell growth (c) and invasion (d) of indicated Src-transformed NIH3T3 cells. **e**: the level of Src expression and activity in SW620 colon cancer cells transduced with indicated Src constructs. **f, g**: anchorage independent growth (f) and invasion (g) of SW620 cancer cells expressing indicated Src constructs. The histograms show the percentage of colonies in soft agar normalized to the maximal condition set a 100% (colonies) and the percentage of migrating cells in the matrigel matrix normalized to control condition set at 100% (cell invasion). Is shown the mean ± SEM; n>3; ns: p>0.05; *p<0.05; **p<0.01; p<0.001; Student’s *t* test.

We next addressed the importance of ULBR in Src transforming activity in human cancer. In spite of the fact that *SRC* somatic mutations are rarely detected in human malignancies, aberrant Src activity, resulting from pathological deregulation, is a bad prognosis maker in epithelial tumors and has important roles during tumor development/progression (2,3). In colon cancer cells, regulation of Src signaling is highly perturbed due to defects in the regulation of its catalytic activity due to CSK inactivation (19), which mediates Src-Tyr530 phosphorylation (4). Defects in the regulation of Src substrates degradation, due to inactivation of the inhibitory signaling protein SLAP in these tumor cells also participates in Src oncogenic signaling (20). As a result, ectopic expression of wild type Src in SW620 colon cancer cells, which originate from a lymph node metastasis and express low levels of endogenous Src, strongly increases their growth and invasive and abilities (21,22) (Fig 2e-g). Expression of ULBR mutated Src produced similar results as in fibroblasts, i.e. a strong diminution of both anchorage-independent cell growth and cell invasion (Fig 2f&g). Importantly, subcutaneous injection of these tumor cells in *nude* mice produced similar results. In this experimental *in vivo* cancer model, Src expression enhanced tumor development by 8-fold as compared to control cells, while this effect was reduced by 60% upon Src3A mutant expression (Fig 3a &b). Immunohistochemical analysis of tumor sections showed a significant reduction in colon cancer cell proliferation in Src3A samples (Fig 3c). In contrast to wild-type Src, Src3A significantly increased tumor cells apoptosis (Fig 3d). Src also induced a substantial increase in tumor angiogenesis, important for tumor progression. While the overall length of tumor vessels was not significantly modified by ULBR inactivation, we noted a significant diminution of nascent tumor vessels in Src3A expressing tumors, suggesting an implication of this region during tumor angiogenesis (Fig 3e).

**Figure 3.**
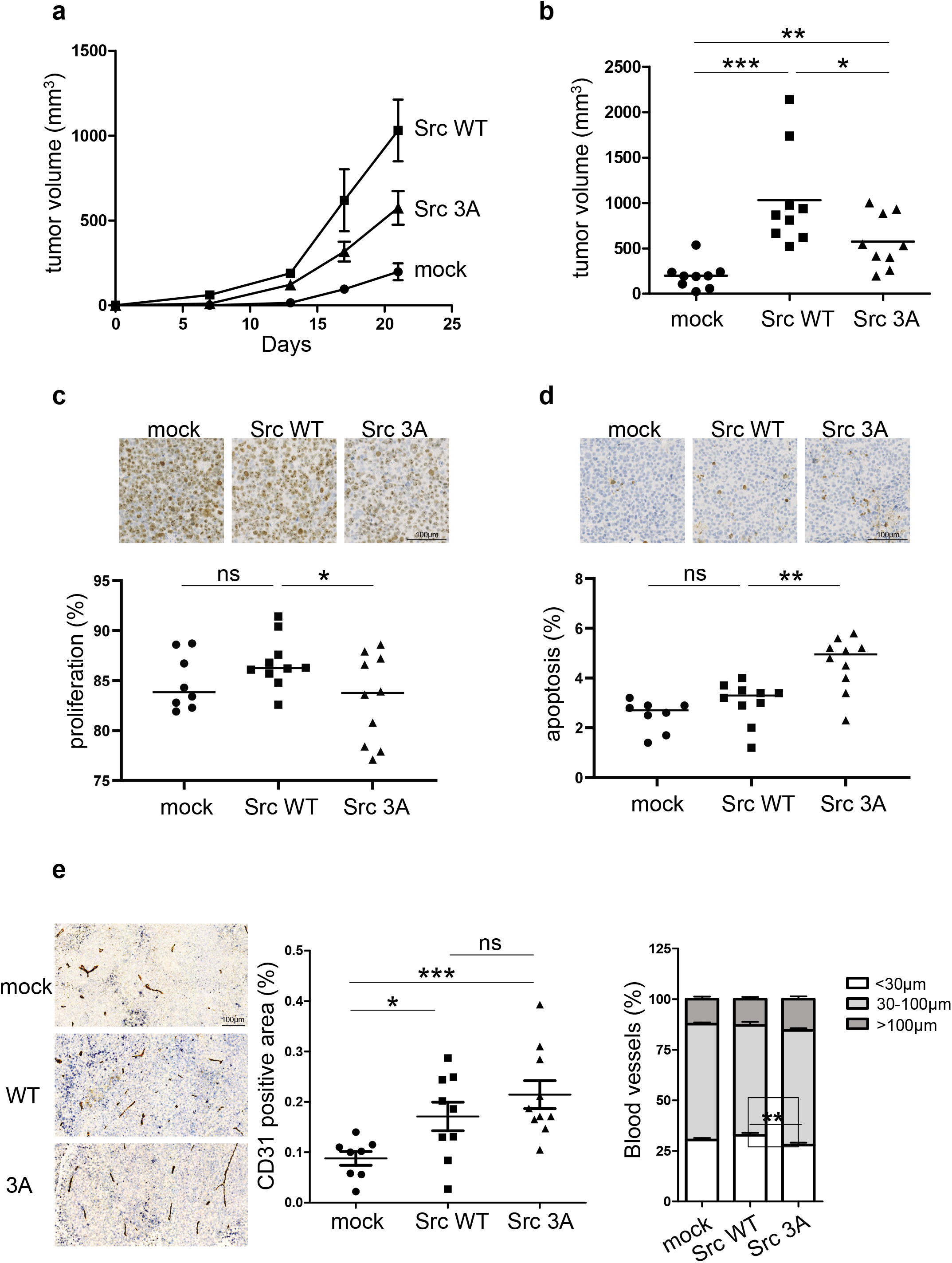
ULBR inactive mutations affects Src tumor activity in nude mice. **a, b**: timecourse of tumor development in nude mice subcutaneously inoculated with SW620 tumor cells that were transduced with control (mock) or indicated Src construct. **c-e**: Analysis of tumor cell proliferation (c), apoptosis (d) and angiogenesis (e) from indicated tumor sections. Representative sections and quantification of immunohistochemical analysis showing tumor cell proliferation (anti-ki67), apoptosis (anti-cleaved Caspase 3) and angiogenesis (anti-CD31; length of blood vessels) in xenograft tumors derived from SW620 cells transduced with indicated Src construct. Is shown the mean ± SEM; n>8 mice per cohort; ns: p>0.05; *p<0.05; **p<0.01; ***p<0.001; Student’s *t* test.

### ULBR inactivation affects Src oncogenic signaling

We next investigated the mechanisms involved in Src-ULBR function. Oncogenic SrcY530F induced a large increase of protein tyrosine phosphorylation in NIH3T3 cells, which was substantially reduced upon ULBR inactivation (Fig S1a). ULBR mutants also showed a reduction in SFKs activity, as measured by the level of their conserved tyrosine phosphorylation localized in the activation loop (i.e. pTyr419 in Src) (Fig S1a), with respect to the one observed in SrcY530F. This suggests that, in the context of oncogenic SrcY530F in NIH3T3 cells, ULBR regulates both substrate phosphorylation and SFK activity. Similar results were observed in HEK293T cells, in which transiently expressed Src resulted in high level of Src activity due to high ectopic kinase expression and low endogenous CSK levels (23). In this context, Src induced a large increase in protein tyrosine phosphorylation, which was dependent upon a functional ULBR (Fig 4a). Src3A also showed a 20% reduction in SFK activity as compared to wild-type Src (Fig 4a). We next searched for a similar mechanism operating in human cancer. Consistent with a robust transforming activity, retroviral transduction of wild-type Src in SW620 cells increased protein tyrosine phosphorylation. As in HEK293T cells, 3A ULBR mutation caused a reduction in substrate phosphorylation but a reduction in active Src level was not evident (Fig 4b). The regulatory role of ULBR on Src signaling was next confirmed on MAPK activity, an important downstream effector of Src transforming activity in epithelial cells (24). ULBR mutations reduced Src-induced p42/44 MAPKs activation both in HEK293T cells and SW620 tumor cells (Fig 4a & b). A similar effect of ULBR inactivation was observed on oncogenic Src signaling in NIH3T3 cells. Src has been shown to induce fibroblasts cell transformation by a p38 MAPK and Stat3-dependent signaling mechanism (25,26). Accordingly, SrcY530F transforming activity was accompanied by an increase in pTyr507-Stat3 and p38 signaling, which was reduced upon ULBR inactivation (Fig S1b).

**Figure 4.**
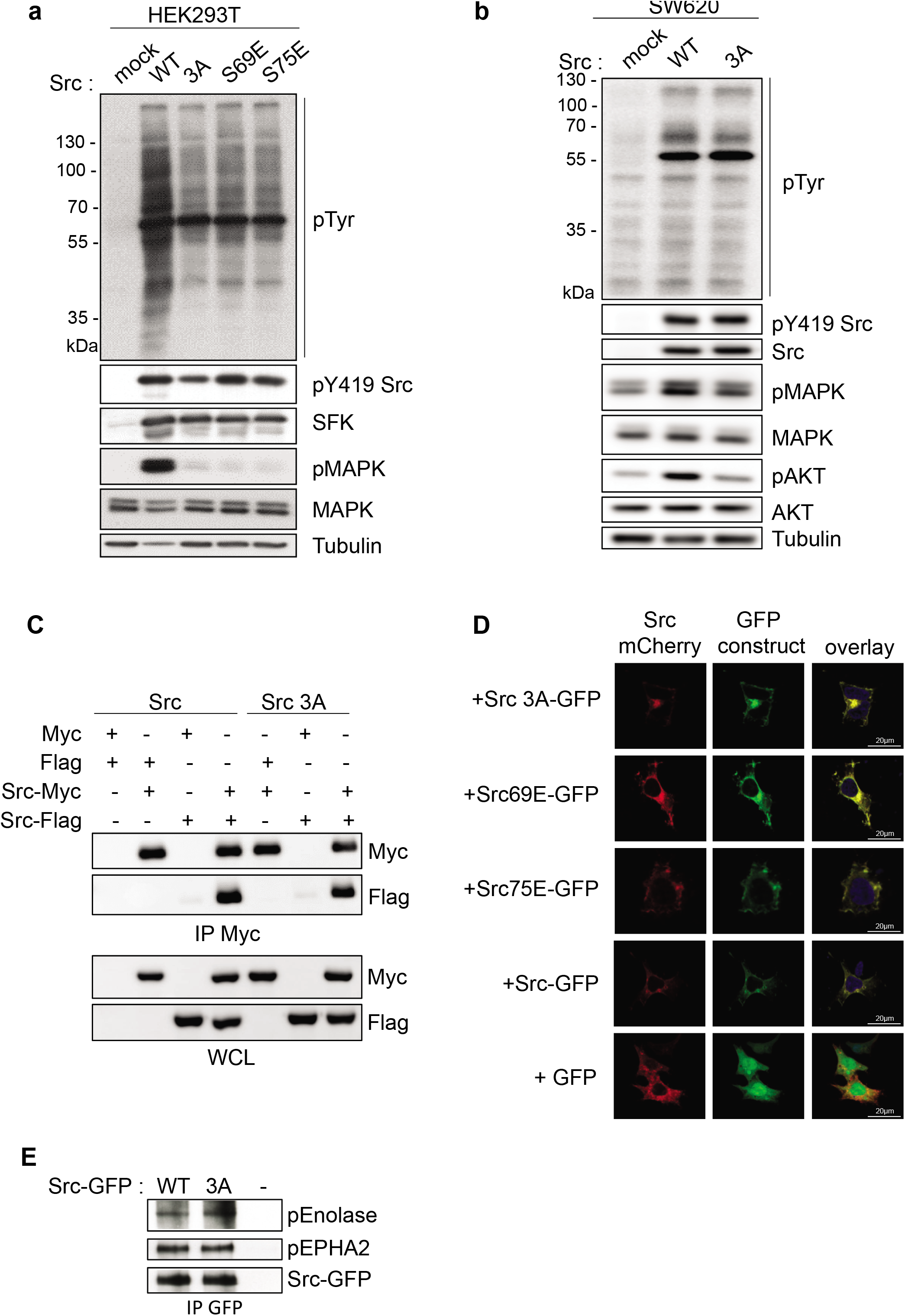
ULBR inactive mutation inhibits MAPK signaling without affecting Src localization and kinase activity. **a, b:** ULBR regulates Src-induced protein tyrosine phosphorylation and of p42/44 MAPK activation. Immunoblots of whole-cell lysates showing cellular protein tyrosine phosphorylation and of p42/44 MAPK activity in SW620 cells (a) and HEK293T cells (b) transduced with indicated Src constructs. **c**: Src dimerization is not affected by ULBR inactivation. HEK293T cells were transfected with the indicated constructs. Src-Myc proteins were immunoprecipitated (IP) from cell lysates and immunoblotted with the indicated antibodies. Immunoblots of whole-cell lysates were also performed as indicated. **d**: Representative confocal image of direct fluorescence of HEK293T cells co-expressing Src-mCherry and indicated Src-GFP ULBR mutants. The overlay is also shown. **e**: In vitro kinase assay of purified Src-GFP and Src3A-GFP that were expressed in HEK293T cells using indicated substrate. The level of immunoprecipitated Src-GFP proteins and tyrosine phosphorylation of indicated substrate is shown.

We next searched for the molecular mechanisms implicated in ULBR-dependent Src substrates phosphorylation. The Src-UD has been suggested to participate in protein dimerization, enabling kinase activation (12,27). We tested the possible ULBR contribution to this molecular process. For this, Src constructs tagged with a hemagglutinin (HA) or a FLAG sequence at the C terminus were co-transfected in HEK293T cells, and Src self-association was assessed by coimmunoprecipitation. Src dimerization was not detected, unless when using stringent lysis conditions (ie RIPA buffer) (19,28). This suggests that dimerization may occur in cholesterol-enriched membrane domains, consistent with lipid binding-induced dimerization/oligomerization observed in SH4 myristoylated Src derivatives (27,29). Using these conditions, we did not detect any effect of ULBR inactivation on Src self-association, indicating that this conserved region may not be involved in kinase dimerization (Fig 4c). We next evaluated the role of ULBR on Src subcellular localization. For this, Src constructs (wildtype or ULBR mutants) were generated with either a GFP or mCherry tag at the C terminus together with a spacer (GluX3) for molecular constraint limitation between GFP (or mCherry) and Src. Wild-type Src fused to mCherry was co-expressed with Src-ULBR mutants fused to GFP in HEK293T cells and their co-localization was analyzed by direct fluorescent microscopy. We found an almost strict co-localization between wild-type Src and ULBR-Src mutants, i.e. at perinuclear membranes, endocytic vesicles and membrane cell periphery, suggesting that ULBR inactivation had no effect on Src sub-cellular localization (Fig 4d). Since the cellular Src kinase activity was affected by ULBR inactivation, we analyzed the impact of ULBR inactivation on Src kinase activity *in vitro* using enolase, or EPHA2 as substrates (21). No difference between purified Src-GFP and Src3A-GFP kinase activity was detected in respect to enolase substrate concentration or kinase duration. (Fig 4e & S2). Similarly, no effect of ULBR inactivation was observed with EPHA2, suggesting that the lack of effect of ULBR mutation *in vitro* is not substrate specific. Altogether, these data indicate that the observed *in vivo* effects with ULBR mutants depend on Src membrane anchoring but not on cell compartmentalization or lipid-induced kinase dimerization.

### Phospho-proteomic analyses of ULBR-Src signaling in tumor cells

We next characterized ULBR-dependent Src phospho-signaling by proteomic methods. SW620 cancer cells were retrovirally transduced with mock (control), wild-type Src and Src3A constructs. A global tyrosine phospho-proteomic analysis was first performed by phosphotyrosine peptide immune-purification from trypsin-digested cell lysates followed by label-free mass spectrometry-based quantification (30). From this analysis, we detected 279 phosphopeptides in control cells with a log2 fold change (FC) ≥1 upon Src (or Src3A) expression (table S1). These Src substrates were essentially composed of signaling proteins, regulators of cell adhesion, trafficking, mRNA maturation, protein synthesis and cell metabolism (table S1), consistent with previous studies (21,22). We next examined how these peptides distribute into wild type Src and Src3A expressing samples, using a log2 FC ≥ 2 threshold. Src- and Src3A-induced protein tyrosine phosphorylation showed very similar profiles (Fig 5a and table S1), indicating that ULBR has no major impact on Src substrate phosphorylation and/or this region may regulate subtle phosphorylation changes that could not be detected from the label free MS analysis We next profiled ULBR signaling by probing a phospho-receptor tyrosine kinase (RTK) antibody array (Fig 5b & S3). This biochemical survey revealed that Src has a large impact on tyrosine phosphorylation of RTKs (>2-fold increase for 13/49 RTKs), including adhesive receptors DDR2 and EPHs, growth factors receptors EGFR, MET, FGFR3 and AXL an RYK, a co-receptor of the Wnt/beta-catenin signaling. Interestingly, Src-ULBR had a modulatory role on Src-induced RTKs phosphorylation, i.e. a positive effect on EGFR, MET, FGFR3, RYK and EPHA2 phosphorylation and an inhibitory effect on EPHA5, EPHB2, B4 and B6 phosphorylation (Fig 5b, S3a & b). We also complemented this analysis by probing a phospho-kinase antibody array (Fig 5c), which revealed that Src expression activates Stats (2, 5 and 6) and SFKs (Fyn, Yes, Lck, Lyn and Hck) signaling proteins. Interestingly, while Src-ULBR was dispensable for most of the probed phospho-signaling activities, it was required for Src-induced Fyn activation (i.e. pTyr420-Fyn level) and activation of the downstream effectors p42/44 MAPKs (i.e. pThr202/pTyr204 level) in these cancer cells (Fig 5c & S3b). We thus concluded that ULBR modulates tyrosine phosphorylation of specific membrane-localized substrates, e.g. RTKs and SFKs.

**Figure 5.**
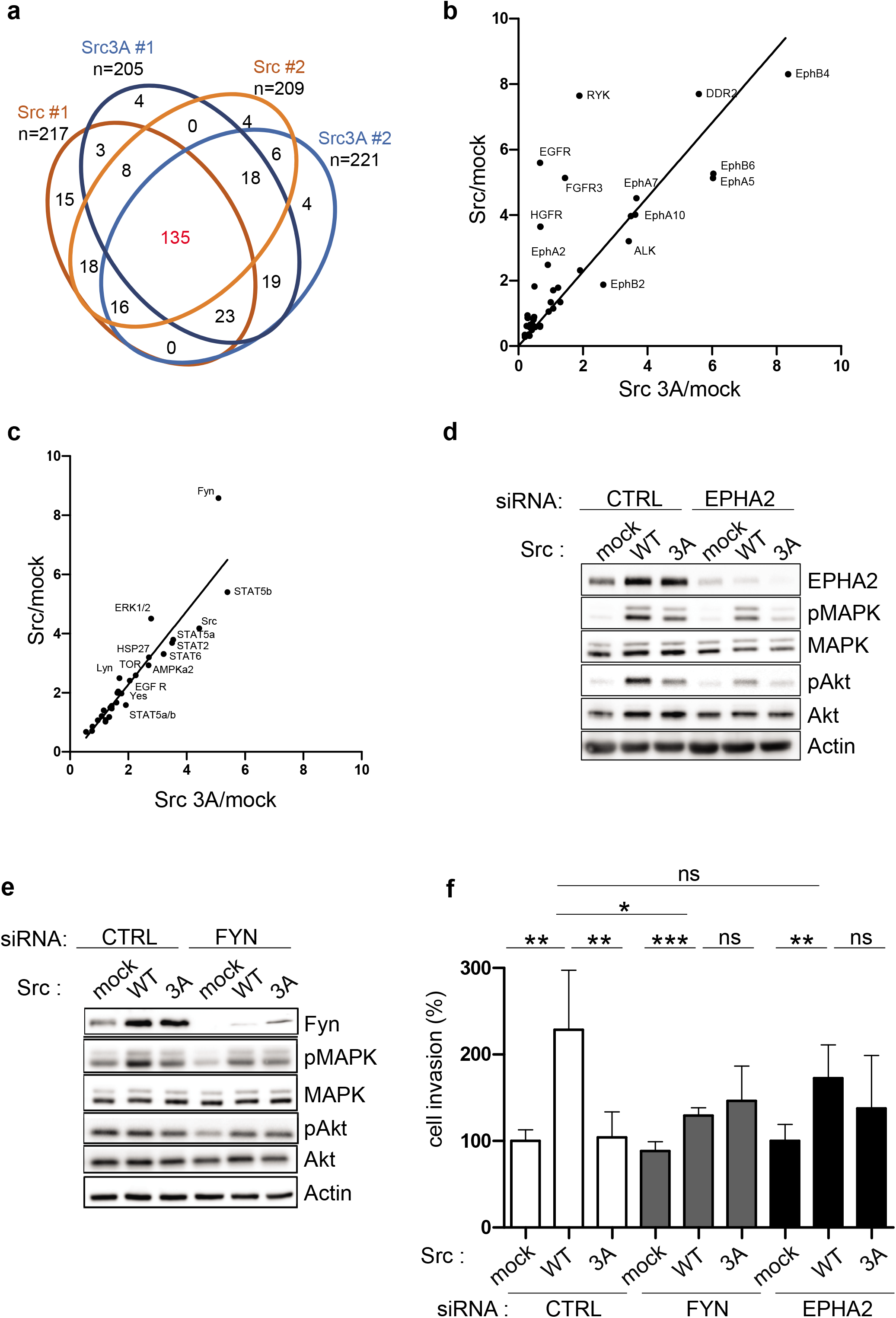
phospho-proteomic analysis of Src-ULBR signaling in SW620 cancer cells. **a:** A label-free quantitative phospho-proteomic analysis centered on tyrosine phosphorylation. A Veen diagram where quantified phospho-peptides were sorted as differentially phosphorylated from the control condition (mock) (log2FC ≥1) in the indicated Src (or Src3A) conditions. **b**: a phospho-RTK array approach. Comparison of Src and Src3A induced tyrosine phosphorylation of RTKs as revealed by phospho-RTKs array. Is shown the relative level of indicated tyrosine phosphorylated RTKs from mock condition. **c**: a phospho-kinase array approach. Comparison of Src and Src3A induced phospho-signaling as revealed by a phospho-signaling kinase array. Is shown the relative level of indicated phosphorylated signaling proteins from mock condition (SW620 cells infected with control retroviruses) **d-f**: Fyn and EPHA2 are important mediators of ULBR-Src signaling in SW620 cancer cells. **d, e**: biochemical analysis of p42/44 MAPK and Akt activity in SW620 expressing or not Src or Src3A mutant as shown and transfected with indicated siRNA. The level of EPHA2, pY594-EPHA2 (pEPHA2) and Fyn is also shown. **f:** cell invasion of SW620 expressing or not Src or Src3A mutant and transfected with indicated siRNA. The histograms show the percentage of migrating cells in the matrigel matrix normalized to control condition set at 100% (cell invasion). Is shown the mean ± SEM; n>3; ns: p>0.05; *p<0.05; **p<0.01; p<0.001; Student’s *t* test.

### EPHA2 and Fyn as important mediators of Src-ULBR signaling in tumor cells

Finally, we aimed at validating these results functionally. ULBR inactivation reduced Src-induced p42/44 MAPK and Akt signaling leading to SW620 cell invasion (Fig 2f and 4b) and we interrogate the role of two ULBR-dependent Src substrates on this signaling response, i.e. EPHA2 and Fyn. EPHA2 is aberrantly stabilized in colon cancer cells by a Src-dependent mechanism implicating inactivation of the SLAP-UBE4A ubiquitination complex (20). Moreover, EPHA2 phosphorylation on Tyr594 by Src amplifies Akt signaling, leading to colon cancer cell invasion (20). Consistent with our proteomic analyses, Src-ULBR regulated EPHA2-Tyr594 phosphorylation (Fig S3a). At the functional level, *EPHA2* silencing inhibited Src signaling leading to Akt and p42/44 MAPK activation and cell invasion (Fig 5d, f and S4); however, it had no clear effect on Src3A signaling responses. EPHA2 is thus an important mediator of ULBR-Src signaling in SW620 cells. The fact that Src induces activation of other SFKs, e.g. Fyn, in colon cancer cells raises the idea that Src interacts with other SFKs to induce oncogenic signaling. Consistently, Src expression increased Fyn protein levels (Fig 5e), likely via a posttranscriptional mechanism because it had no impact on *FYN* transcript level (Fig S4b). Functionally, Fyn depletion specifically inhibited Src-induced p42/44 MAPK activation and cell invasion (Fig 5f and S4). However, no such inhibitory effect was observed on Src3A signaling placing Fyn as an additional effector of Src-ULBR signaling. The interaction between Src and Fyn signaling was next confirmed by showing a potentiating effect of Fyn on Src-induced p42/44 MAPK activation in HEK293T cells (Fig S4d). However, this signaling response was reduced upon Src-ULBR mutation, supporting further a role for Src-UD in Src-Fyn signaling (Fig S4d). Overall, our results point to an important role of Src-ULBR in the regulation of membrane substrates phosphorylation, essential for Src tumor signaling.

## DISCUSSION

Most eukaryotic proteins have intrinsically disordered regions (IDRs) that challenge the classical structure-function paradigm. Here, we uncover an important role of this molecular property made by Src-UD in cancer development. The biological roles of IDRs contained in SFK have remained unclear, despite strong insight into Src regulation. Previous mutagenesis experiments centered in this region did not reveal any clear oncogenic activity (7), possibly because Src-IDR may contain opposite regulatory sequences that would not be detected with this approach. Consistent with this idea, our mutagenesis analysis guided from our NMR data led to specifically inactivate the ULBR, a small region conserved in Src-UD, which revealed its essential role in Src tumor activity. This sequence may however not regulate all Src transforming functions, as suggested from Src-induced morphological changes in mouse fibroblasts. Actually, Src-ULBR may act as a fine-tuning mechanism, which may be exacerbated upon Src overactivation to promote cancer development. This fine-tuning mechanism is supported by previous works on the Src capacity to induce Xenopus oocytes maturation (10), Src regulation of retinal ganglion cell survival or postmitotic neuron function by Ser75 phosphorylation (31,32) and is consistent with a recent Src optogenetic study (33).

Our study also brings valuable molecular insight into the mechanism of Src regulation by its unstructured region. Previous studies reported that Lck-IDR can mediate protein interaction by adopting an organized structure (34). Whether IDR of other SFKs display similar molecular property is not known, although Src-UD was involved in protein interactions (35). IDRs adopt multiple conformations that are sensitive to the environment and, through multiple weak interactions in a fuzzy complex, may direct the activity of folded domains of signaling proteins towards different pathways (8). Our results support such a Src-IDR regulatory mechanism at the plasma membrane since ULBR regulates Src membrane anchoring and phosphorylation of essential membrane localized substrates of tumor signaling, notably RTKs and SFKs. An unsuspected finding from this work is the Src capacity to activate additional SFKs, such as Fyn, to promote oncogenic signaling. This result suggests the existence of a SFK network involved in cancer development and uncovers a novel layer of Src signaling complexity, which deserves further investigation.

By focusing on ULBR, this work started addressing the biological role of Src-IDR but existing molecular studies uncovered additional regions involved in the fuzzy complex made by SFKs-IDR, which would also contribute to Src regulation. Unravelling the biological role of these regions may bring a more complete view on the complexity of Src regulation by its IDR. Finally, Src has been identified as an attractive target in oncology but Src inhibitors developed for the clinic gave disappointing results in colon cancer, probably because of high toxicity and inefficient Src signaling inhibition (3). Interestingly, structural analyses of non-catalytic domains of TKs have revealed unique modes of kinase regulation (36), which resulted in the development of allosteric inhibitors with improved anti-tumor activities, as reported for asciminib in chronic myeloid leukemia (37). We thus propose that targeting the IDR fuzzy complex with small molecules would circumvent some of these issues and therefore may define an attractive strategy to block Src tumor activity in human cancer.

## MATERIAL AND METHODS

### Antibodies

anti-p42/44 MAPKs, anti-p42/44 MAPKs pT202/Y204, anti-p38 MAPK, anti-p38 MAPK pT180/Y182, anti-AKT, anti-AKT pS473, anti-EPHA2 pY594, anti-Src pY418, anti-Stat3 pY705, anti-Cyclin D3 (Cell Signaling Technology), anti-EPHA2, anti-Stat3, anti-Myc (Santa Cruz Biotechnology), anti-Src specific (2.17) antibody was a generous gift from Dr S Parsons (University of Virginia, VA, USA), anti-pTyr clone pY1000 Sepharose bead conjugated (PTM Scan, Cell Signaling Technology); anti-FLAG (Sigma Aldrich), and anti-GFP (Chromotek), anti-tubulin (gift from N. Morin, CRBM, Montpellier, France); anti-pTyr 4G10 (gift from P. Mangeat, CRBM, Montpellier, France); anti-cst1 (that recognizes Src, Fyn and Yes) was described in (38). Anti-rabbit IgG-HRP and anti-mouse IgG-HRP (GE Healthcare). Anti-Mib1h (Dako)(KI67), anti-active-caspase3 (AP175; cell signaling), anti-CD31 (Ab28364, Abcam).

### Phylogenetic analyses

Nucleic and protein SFK sequences were retrieved from NCBI annotated nr database (http://www.ncbi.nlm.nih.gov). Accessions are listed Table S1. Protein sequences were aligned using MAAFT v7.450 (39). Nucleic sequence alignments were based on protein alignments. Phylogenetic trees were estimated by PhyML (40), using the General Time Reversible (GTR) model with invariant and gamma decorations. Nonsynonymous versus synonymous substitution ratios (ω= dN/dS) were calculated using PAML 4.4 (41). We compared the “one-ratio” model (a single ω ratio for the entire tree) and the “two-ratio” model (distinct ω values for each of the Src, Fyn and Yes branches) by using the likelihood ratio test.

### Plasmids

pMX-pS-CESAR retroviral vector expressing human Src was described in (20). pMX-Src L63A/F54A/G65A (Src 3A) was described in (11). The other plasmids were obtained by PCR using the QuickChange Site-Directed Mutagenesis Kit (Stratagene) using specific oligonucleotides as follows: Src S69E, Forward-5’CGGAGGCTTCAACGCCTCGGACACCGT3’, Reverse-5’ACGGTGTCCGAGGCGTTGAAGCCTCCG3’; Src S75E, Forward-5’GACACCGTCACCGCCCCGCAGAGGG3’, Reverse-5’CCCTCTGCGGGGCG GTGACGGTGTC3’; Src Y530F (Src YF), Forward-5’CGGGCTGGAACTGGGGCTCGGTG G3’, Reverse-5’CCACCGAGCCCCAGTTCCAGCCCG3’. Each of the Y530F counterparts Src L63A/F54A/G65A/Y530F (SrcYF 3A), Src S69E/Y530F (SrcYF S69E); Src S75E/Y530F (SrcYF S75E) were obtained by adding the Y530F mutation using the following oligonucleotides Forward-5’ TCTCGAGCTCAAGCTTAGTACCCTTCACCATGGGTAGC AACAA3’, Reverse-5’ GGCGACCGGTGGATCCGAGCCGGAGCCGAGGTTCTCCCCGGGCTGGTA3’. Src-GFP and Src-mCherry constructs were obtained by insertion of Src sequence (or Src mutants) in pEGFP-N1 and pmCherry N1 respectively including a GluX3 spacer. Src-Flag and Src-Myc constructs were obtained by inserting Src and Src3A sequences in pcDNA3 vectors. pSG5 Fyn construct was described in (18).

### Cell cultures, retroviral infections and transfections

Cell lines (NIH3T3, HEK293T and SW620 cells) (ATCC, Rockville, MD) were cultured, transfected and infected as described in (20). Stable cell lines were obtained by fluorescence-activated cell sorting. For siRNA transfection, 2.10^5^ cells were seeded in 6-well plates and transfected with 20 nmol of siRNA and 9 μl of Lipofectamine RNAi Max according to the manufacturer’s protocol (ThermoFisher Scientific). A scramble siRNA (siMock) 5’TTCTCCGAACGTGTCACGTTT3’ was used as a negative control (Eurofins). The following siRNAs were used for functional assays: siRNA FYN#1 (Cell Signaling Technology #12473), siRNA FYN#2 5’GGCCCTTTATGACTATGAATT3’, siRNA EPHA2#1 5’GCAGT ATACGGAGCACTTCTT3’, siRNA EPHA2#2 5’GTATCTTCATTGAGCTCAATT3’ (Eurofins).

### Biochemistry

Immunoprecipitation and immunoblotting were performed as described in (20). Kinase assays were performed as described in (20) using 200ng of purified EPHA2 recombinant protein (OriGen Technologies) or indicated concentration of purified Enolase (Sigma Aldrich), in the absence or presence of about 50ng of purified Src-GFP (or Src3A-GFP as indicated) in the presence of 0.1mM ATP Lithium Salt (Roche Diagnostics) in kinase buffer (20mM Hepes pH6.5, 10mM MnCl_2_, 1mM DTT) for indicated time at 30°C. Src-GFP purification was performed by anti-GFP immunoprecipitation from HEK293T cells transfected with Src-GFP (or Src3A-GFP) construct.

### RNA extraction and RT-quantitative PCR

mRNA was extracted from cell lines and tissue samples using the RNeasy plus mini kit (Qiagen) according to the manufacturer’s instructions. RNA (1μg) was reverse transcribed with the SuperScript VILO cDNA Synthesis Kit (Invitrogen). Quantitative PCR (qPCR) was performed with the SyBR Green Master Mix in a LightCycler 480 (Roche). Expression levels were normalized with the Tubulin human housekeeping gene. Primers used for qPCR: Tubulin, Forward-5’CCGGACAGTGTGGCAACCAGATCGG3’, Reverse-5’TGGCCAAAAGGACCTGAGCG AACGG3’; Fyn, Forward-5’TGACCTCCATCCCCAACTA3’, Reverse-5’TTCCCACCAATCTCCTTCC3’; EPHA2, Forward-5’GGGACCTGATGCAGAACATC3’, Reverse-5’AGTTGGTGCGGAGCCAGT3’.

### Cell imaging

HEK293T cells plated on glass coverslips coated with fibronectin were transfected with Src-GFP and Src-mCherry constructs for 24 hours and subcellular Src distribution was analyzed after cell fixation (4% paraformaldehyde) by direct fluorescence using confocal microscopy. Src-transformed NIH3T3 cells were plated on glass coverslip coated with fibronectin for 24 hours and actin was visualized with Texas red-conjugated phalloidin (1:200 dilution) after cell fixation (4% paraformaldehyde) and permeabilization (0.05% TRITON for 10 min at room temperature).

### Soft agar colony formation and cell invasion assay

Colonies formation: 1 000 cells per well were seeded in 12-well plates in 1ml DMEM containing 10% FCS and 0.33% agar on a layer of 1ml of the same medium containing 0.7% agar. After 18-21 days, colonies with > 50 cells were scored as positive. Cell invasion assay was performed as described in (30) using Fluoroblok invasion chambers (BD Bioscience) in the presence of 100 μl of 1-1.2 mg/ml Matrigel (BD Bioscience).

### Phospho-proteomic analyses

Quantitative phosphoproteomics was performed as in (30). Briefly, SW620 cells were lysed in urea buffer (8M urea in 200mM ammonium bicarbonate pH 7.5). Phosphopeptides were purified after tryptic digestion of 20 mg (for cells) or of 35 mg (for mouse tumors) total proteins using the PTMScan^®^ Phospho-Tyrosine Rabbit mAb (P-Tyr-1000) Kit (Cell Signaling Technology), according to manufacturer’s protocol. An additional enrichment step using the IMAC-Select Affinity Gel (Sigma Aldrich) was performed to increase the phosphopeptide enrichment. Purified phosphopeptides were resuspended in 10% formic acid and two technical replicates for each sample were analyzed. Phospho-kinase arrays: proteome profiler human phospho-kinase array and human phospho-RTK array kits including 49 RTKs were purchased from R&D Systems. Indicated SW620 cells were lysed, and 300 μg of protein lysates were subjected to western blotting according to the manufacturer’s protocol. Signals on membranes were quantified using the Amersham Imager 600 (GE Healthcare) from 2 independent biological replicates.

### *In vivo* experiments and Immunohistochemistry (IHC)

*In vivo* experiments were performed in compliance with the French guidelines for experimental animal studies (Direction des services vétérinaires, ministère de l’agriculture, agreement B 34-172-27). 2×10^6^ SW620 cells (or derivatives) were subcutaneously injected in the flank of 5-week-old female athymic nude mice (Envigo). Tumor volumes were measured as the indicated intervals using calipers. After 24 days, tumors were excised, weighed and cryopreserved or processed for subsequent immunohistochemistry analysis as described in (22).

### Statistical analysis

All analyses were performed using GraphPad Prism. Data are presented as the mean ± SEM. When distribution was normal (assessed with the Shapiro Wilk test), the twotailed *t* test was used for between-group comparisons. In the other cases, the Mann-Whitney test was used. Statistical analyses were performed on a minimum of three independent experiments. The statistical significance level is illustrated with p values: *p≤0.05, **p≤0.01, ***p≤0.001.

## ACKNOWLEDGEMENTS

We thank the Montpellier RIO Imaging and RHEM platforms for imaging and immunohistochemistry analyses, and the Proteomic Core Platform of EPFL for proteomic analyses. This work was supported by fundacion Marato TV3, La Ligue Nationale Contre le Cancer (LNCC), Montpellier SIRIC Grant «INCa-DGOS-Inserm 6045», CNRS, and the University of Montpellier. RHEM facility supported by SIRIC Montpellier Cancer Grant INCa_Inserm_DGOS_12553, the european regional development foundation and the occitanian region (FEDER-FSE 2014-2020 Languedoc Roussillon), IBiSA and Ligue contre le cancer for processing our animal tissues and histology technics. ML is supported by the Fondation pour la Recherche Médicale (FRM) and the Fondation de France. SR is an INSERM investigator.

## AUTHORS CONTRIBUTION

All authors contributed extensively to the work presented in this paper. Experimental analysis and Data acquisition: EA, ML, MB, VS, BR, EF, FA, RH, JP and AS. Bioinformatics analysis: PF, MA and RH. Writing of the paper: SR and MP. Project supervision: SR.

