## supplementary information for "Regulation of Src tumor activity by its N-terminal intrinsically disordered region"

**Table S1. Label-free tyrosine phospho-proteomic analysis of Src-ULBR signaling in SW620 cells.**

**Figure S1. ULBR signaling in Src-transformed NIH3T3 cells. a**: ULBR regulates SFK activity and protein tyrosine phosphorylation. **b**: ULBR regulates p38 MAPK and Stat3 signaling. **c:** ULBR does not affect Src-induced dissolution of actin fibers. A representative example of actin immunostaining (left) and the quantification (right) of cells transformed with indicated Src mutants showing remaining actin fibres. Is shown the mean ± SEM; n>3; ns: p>0.05; *p<0.05; **p<0.01; p<0.001; Student's *t* test.

**Figure S2. ULBR does not regulates Src kinase activity in vitro.** Western blotting (**a**) and quantification (**b**) showing the time course and concentration-dependence of Enolase phosphorylation by immunopurified Src-GFP and Src3A-GFP using an in vitro kinase assay described in methods.

**Figure S3. ULBR modulates Src-induced RTK and Fyn tyrosine phosphorylation. a:** ULBR-dependent Src signaling in SW620 cells. **b**: phospho-RTKs (left) and phospho-kinase antibody array analysis (right) using SW620 cells expressing indicated constructs. Specific Src-ULBR substrates are highlighted.

**Figure S4. Fyn and EPHA2 are important mediators of ULBR-Src signaling in SW620 cancer cells. a**: cell invasion of SW620 expressing or not Src or Src3A mutant and transfected with indicated siRNA. **b**: Src does not affect Fyn mRNA level in SW620 cells. Is shown the relative mRNA Fyn level in SW620 cells expressing indicated Src construct and transfected with indicated siRNA. **c**: EPHA2 protein level in SW620 cells expressing indicated Src construct and transfected with indicated siRNA. **d**: ULBR-dependent interaction between Src and Fyn signaling in HEK293T cells.

Figure S1

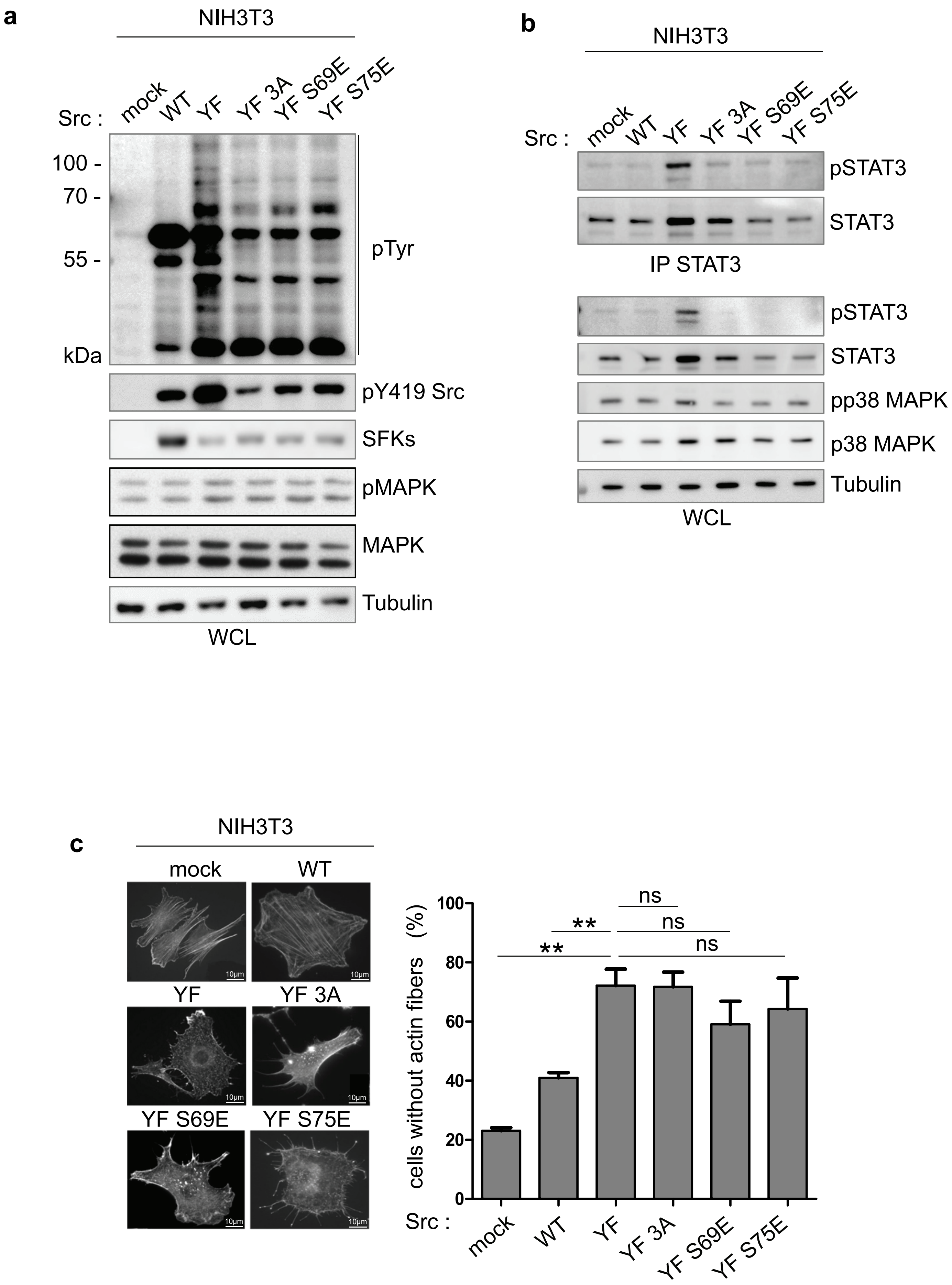
a
NIH3T3
Src :
mock
WT
YF
YF 3A
YF S69E
YF S75E
100 -
70 -
55 -
kDa
pTyr
pY419 Src
SFKs
pMAPK
MAPK
Tubulin
WCL
b
NIH3T3
Src :
mock
WT
YF
YF 3A
YF S69E
YF S75E
pSTAT3
STAT3
IP STAT3
pSTAT3
STAT3
pp38 MAPK
p38 MAPK
Tubulin
WCL
c
NIH3T3
mock
WT
YF
YF 3A
YF S69E
YF S75E
10µm
cells without actin fibers (%)
0
20
40
60
80
100
**
**
ns
ns
ns
Src :
mock
WT
YF
YF 3A
YF S69E
YF S75E

Figure S2

**a**

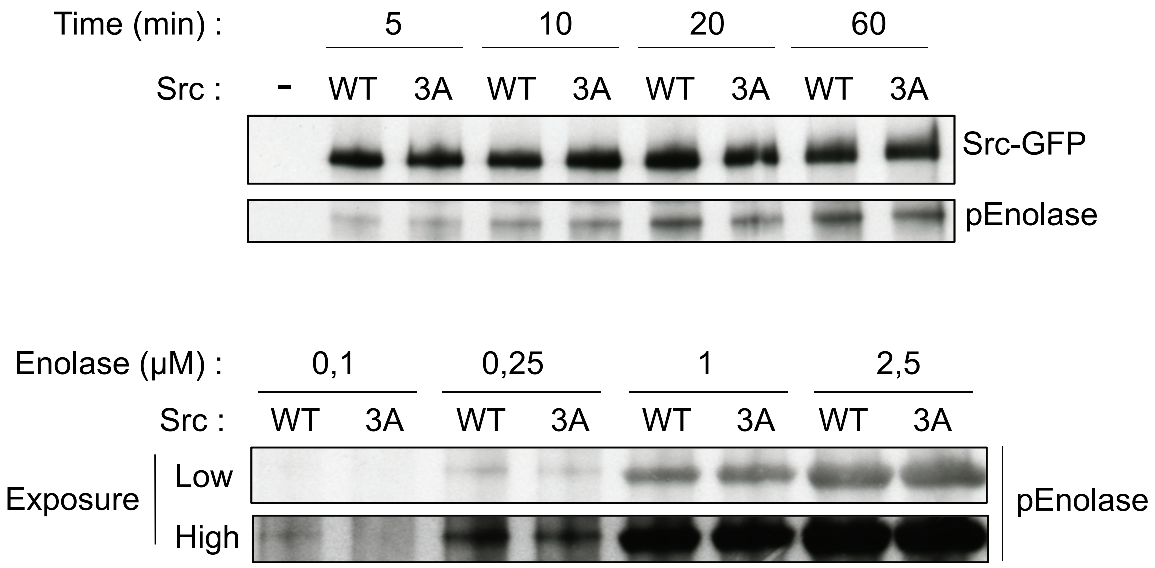
Time (min) :
5
10
20
60
Src :
-
WT
3A
WT
3A
WT
3A
WT
3A
Src-GFP
pEnolase
Enolase (μM) :
0,1
0,25
1
2,5
Src :
WT
3A
WT
3A
WT
3A
WT
3A
Exposure
Low
High
pEnolase

**b**

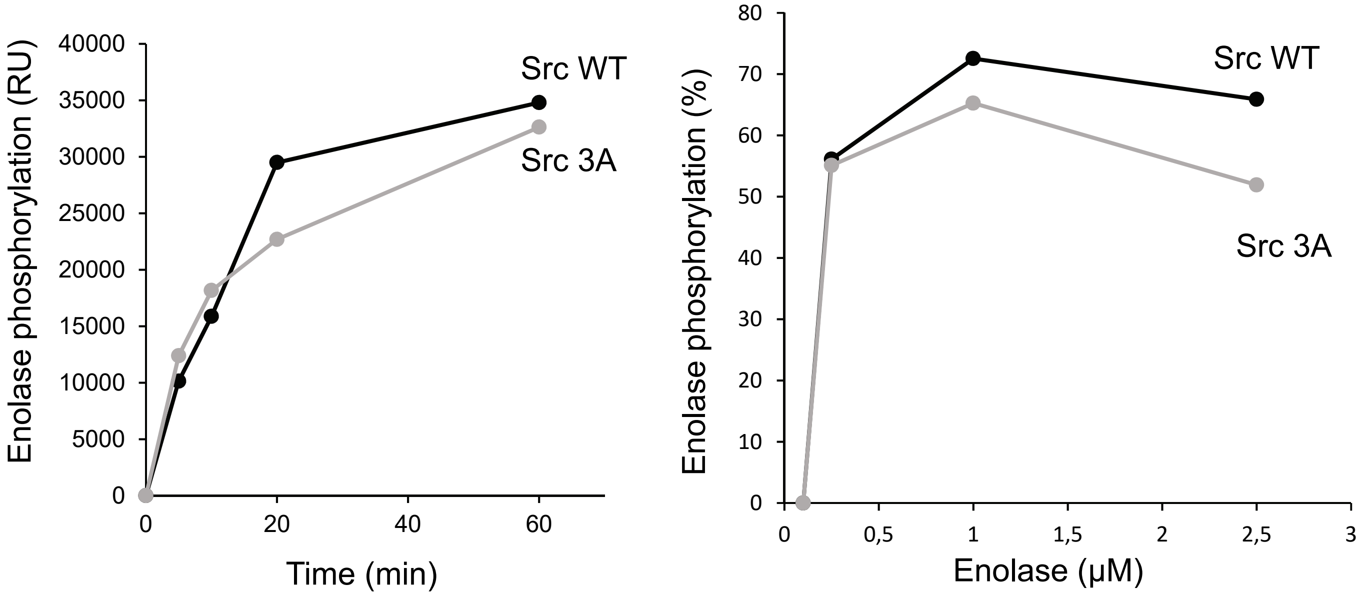
Enolase phosphorylation (RU)
40000
35000
30000
25000
20000
15000
10000
5000
0
0
20
40
60
Time (min)
Src WT
Src 3A
Enolase phosphorylation (%)
80
70
60
50
40
30
20
10
0
0
0,5
1
1,5
2
2,5
3
Enolase (μM)
Src WT
Src 3A

Figure S3

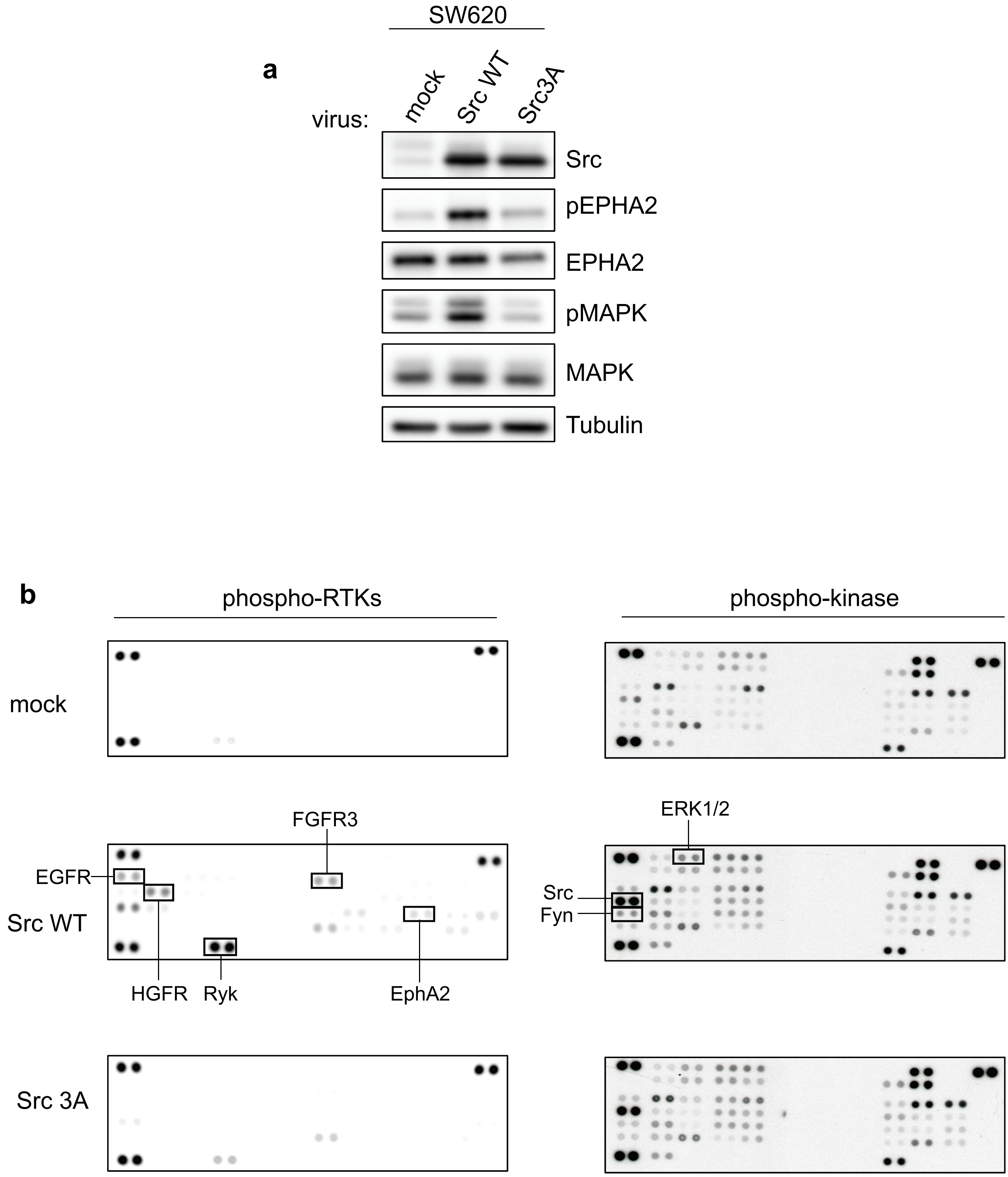

SW620
a
virus:
mock
Src WT
Src3A
Src
pEPHA2
EPHA2
pMAPK
MAPK
Tubulin
b
phospho-RTKs
phospho-kinase
mock
Src WT
Src 3A
EGFR
HGFR
Ryk
FGFR3
EphA2
ERK1/2
Src
Fyn

Figure S4

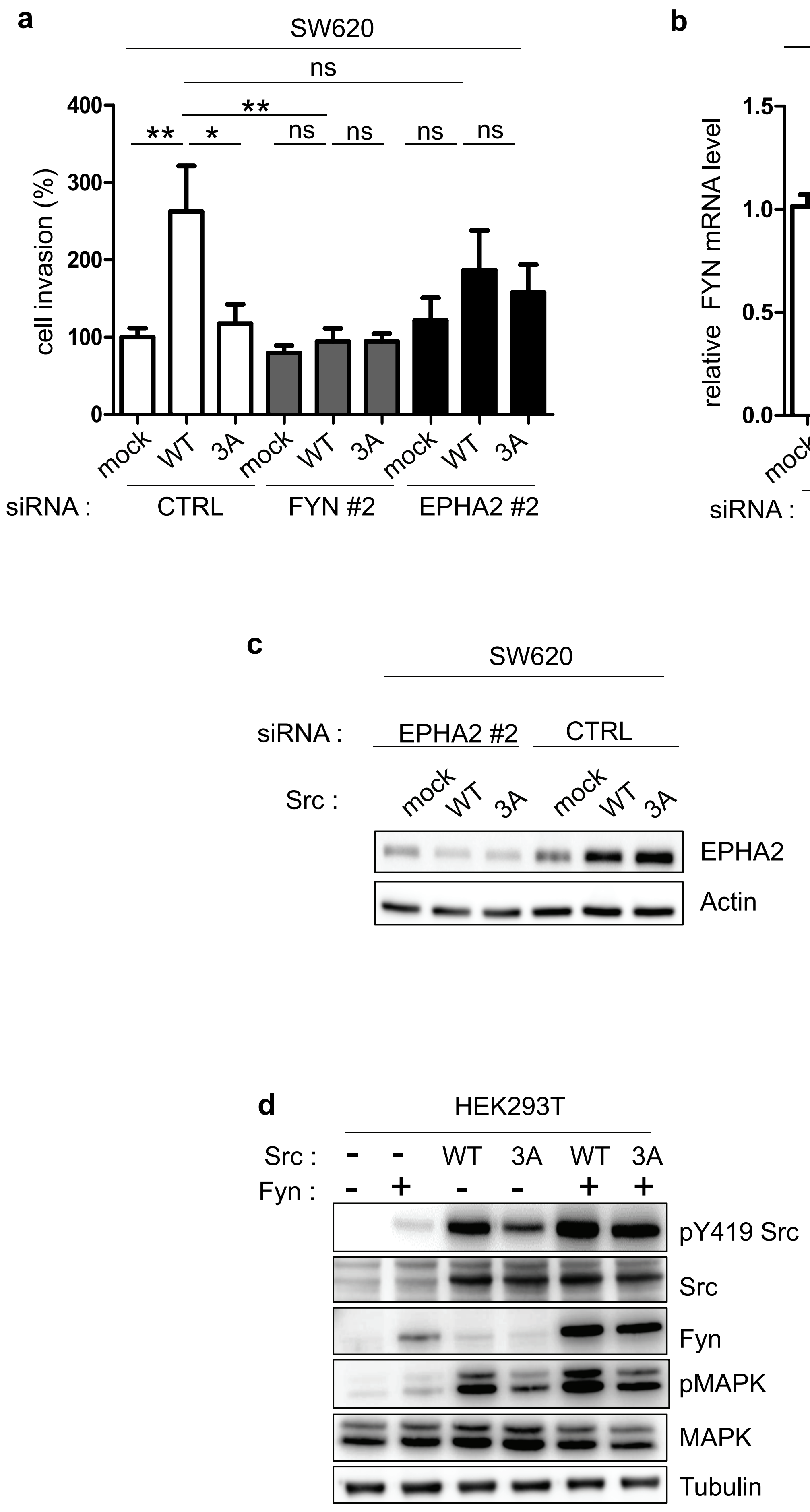
a
SW620
ns
**
**
*
ns
ns
ns
ns
400
300
200
100
0
cell invasion (%)
mock
WT
3A
mock
WT
3A
mock
WT
3A
siRNA :
CTRL
FYN #2
EPHA2 #2
b
SW620
1.5
1.0
0.5
0.0
relative FYN mRNA level
***
***
***
mock
WT
3A
mock
WT
3A
siRNA :
CTRL
FYN #2
c
SW620
siRNA :
EPHA2 #2
CTRL
Src :
mock
WT
3A
mock
WT
3A
EPHA2
Actin
d
HEK293T
Src :
-
-
WT
3A
WT
3A
Fyn :
-
+
-
-
+
+
pY419 Src
Src
Fyn
pMAPK
MAPK
Tubulin

| T..Proteins | T..Positions.within.proteins | T..Leading.proteins | T..Protein | T..Protein.names | T..Gene.names | T..Sequence.window | T..Fasta.headers | T..id | M..Number.of.Phospho..STY. | log Ratios.SW620_3A_01vsSW620_Ctrl_01 | log Ratios.SW620_3A_02vsSW620_Ctrl_02 | log Ratios.SW620_WT_01vsSW620_Ctrl_01 | log Ratios.SW620_WT_02vsSW620_Ctrl_0 | Class | class code |
|---|---|---|---|---|---|---|---|---|---|---|---|---|---|---|---|
| Q5VST6;Q5VST6-2 | 58;58 | Q5VST6 | Q5VST6 | Alpha/beta hydrola | ABHD17B | GSRWTLHLSERAD | >sp\|Q5VST6\|AB17B_ | 3126 | 1 | 4,0355 | 3,2931 | 2,77 | 2,3878 | 4 | **4** |
| P0DMV8-2;P0DMV! | 470;525;52 | P0DMV8-2 | P0DMV8-2 | Heat shock 70 kDa | HSPA1L | LSKEEIERMVQEA | >sp\|P0DMV8-2\|HS71 | 1091 | 1 | 3,3696 | 3,2057 | 4,18 | 2,5043 | 4 | **3** |
| H0Y4R1;P12268 | 386;430 | H0Y4R1 | H0Y4R1 | Inosine-5'-monoph | IMPDH2 | GSLDAMDKHLSS( | >tr\|H0Y4R1\|H0Y4R1_ | 1758 | 1 | 2,3074 | 2,8868 | 2,88 | 3,4163 | 4 | **A** |
| O15357;O15357-2; | 1135;893;1 | O15357 | O15357 | Phosphatidylinositc | INPPL1 | SCSVLQMAKTLSE | >sp\|O15357\|SHIP2_H | 2070 | 1 | 3,5196 | 2,7612 | 3,39 | 3,7265 | 4 | **w** |
| A0A0U1RQL3;O603 | 359;1631;1 | A0A0U1RC | A0A0U1RQL3 | Kinesin-like protein | KIF1B;KIF1Bbeta | PEFEQFQIVPAVE | >tr\|A0A0U1RQL3\|A0 | 890 | 1;2 | 6,417 | 5,5701 | 3,13 | 4,4947 | 4 | **u** |
| P04083 | 207 | P04083 | P04083 | Annexin A1 | ANXA1 | FGVNEDLADSDAI | >sp\|P04083\|ANXA1_ | 2220 | 1 | 2,0149 | 2,3502 | 2,67 | 2,1957 | 4 | **R2** |
| P11279;P11279-2 | 414;361 | P11279 | P11279 | Lysosome-associate | LAMP1 | VLIAYLVGRKRSH | >sp\|P11279\|LAMP1_ | 2347 | 1 | 4,1369 | 4,2101 | 4,62 | 4,1 | 4 | **R1** |
| H0YCG2;P13473-2 | 252;404 | H0YCG2 | H0YCG2 | | LAMP2 | IIVIVIAYVIGRRKS | >tr\|H0YCG2\|H0YCG2 | 1780 | 1 | 2,9264 | 4,3646 | 2,92 | 4,1968 | 4 | **E** |
| Q6IAA8 | 138 | Q6IAA8 | Q6IAA8 | Ragulator complex | LAMTOR1 | PIPFSDLQQVSRIA | >sp\|Q6IAA8\|LTOR1_I | 3143 | 1;2 | 4,9747 | 4,0815 | 5,19 | 4,6395 | 4 | **Total** |
| Q14739 | 595 | Q14739 | Q14739 | Lamin-B receptor | LBR | LVHREARDEYHCK | >sp\|Q14739\|LBR_HU | 3017 | 1 | 2,5184 | 2,8829 | 3,51 | 2,9768 | 4 | |
| P00338;P00338-4;P | 10;10;39;1 | P00338 | P00338 | L-lactate dehydroge | LDHA | _______MATLKDC | >sp\|P00338\|LDHA_H | 2191 | 1 | 1,0429 | 1,2335 | 2,01 | 1,212 | 4 | |
| Q8WWI1-3;E9PMS( | 223;132;17 | Q8WWI1-3 | Q8WWI1-3 | LIM domain only pr | LMO7 | LEDSSFLKRSGRDS | >sp\|Q8WWI1-3\|LMO | 3286 | 1 | 3,9643 | 4,5624 | 3,91 | 4,079 | 4 | |
| Q6ZN28 | 520 | Q6ZN28 | Q6ZN28 | Metastasis-associat | MACC1 | PDPTPNLKRLSNLI | >sp\|Q6ZN28\|MACC1_ | 3163 | 1 | 2,32 | 3,0237 | 2,81 | 3,536 | 4 | |
| H0YBV6;Q9BXY0 | 36;33 | H0YBV6 | H0YBV6 | Protein MAK16 hon | MAK16 | SFKIRTKTQSFCRN | >tr\|H0YBV6\|H0YBV6_ | 1777 | 1 | 2,1003 | 4,1839 | 2,92 | 4,1709 | 4 | |
| Q9Y316-2;Q9Y316;( | 187;210;21 | Q9Y316-2 | Q9Y316-2 | Protein MEMO1 | MEMO1 | QRFRYSYYDESQG | >sp\|Q9Y316-2\|MEM( | 3642 | 1 | 3,2611 | 1,4457 | 3,39 | 2,0557 | 4 | |
| Q9NXC5 | 761 | Q9NXC5 | Q9NXC5 | WD repeat-containi | MIOS | YSCSAVPHQGRGI | >sp\|Q9NXC5\|MIO_HU | 3560 | 1 | 3,3485 | 3,4202 | 3,77 | 3,3625 | 4 | |
| F8WF70;H7C579;Q! | 49;143;49 | F8WF70 | F8WF70 | Omega-amidase NI | NIT2 | QGAKIVSLPECFN! | >tr\|F8WF70\|F8WF70 | 1709 | 1 | 4,2781 | 5,7597 | 3,59 | 5,3175 | 4 | |
| D6RDK6;D6RBN5;Q | 145;172;19 | D6RDK6 | D6RDK6 | OCIA domain-conta | OCIAD1 | PDPNLEESPKRKN | >tr\|D6RDK6\|D6RDK6 | 1468 | 1 | 2,7744 | 3,9019 | 4,61 | 4,7113 | 4 | |
| Q9UQ80;F8VR77;Q | 255;255;20 | Q9UQ80 | Q9UQ80 | Proliferation-associ | PA2G4 | AGQRTTIYKRDPS | >sp\|Q9UQ80\|PA2G4_ | 3627 | 1 | 3,5023 | 4,6368 | 4,43 | 4,7663 | 4 | |
| A0A087WTT1;P119 | 266;291;24 | A0A087W1 | A0A087WTT1 | Polyadenylate-bind | PABPC1;PABPC1L | LKRKFEQMKQDR | >tr\|A0A087WTT1\|A0 | 843 | 1 | 4,1739 | 3,0658 | 4,83 | 2,3142 | 4 | |
| A0A087WTT1;P119 | 272;297;25 | A0A087W1 | A0A087WTT1 | Polyadenylate-bind | PABPC1;PABPC1L | QMKQDRITRYQG | >tr\|A0A087WTT1\|A0 | 844 | 1 | 3,667 | 3,5723 | 3,12 | 2,9121 | 4 | |
| Q8WUM4;Q8WUM | 727;732 | Q8WUM4 | Q8WUM4 | Programmed cell de | PDCD6IP | SIAREPSAPSIPTP | >sp\|Q8WUM4\|PDC6I | 3277 | 1 | 6,8043 | 3,1511 | 6,61 | 2,5254 | 4 | |
| Q9Y446;Q9Y446-2;I | 210;225;15 | Q9Y446 | Q9Y446 | Plakophilin-3 | PKP3 | YSLVSEQLEPAATS | >sp\|Q9Y446\|PKP3_H | 3653 | 1 | 6,3958 | 4,6261 | 6,76 | 5,8657 | 4 | |
| Q8WZ64 | 77 | Q8WZ64 | Q8WZ64 | Arf-GAP with Rho-G | ARAP2 | KQLQIILSKMQDIF | >sp\|Q8WZ64\|ARAP2_ | 3297 | 1 | 5,2057 | 4,55 | 3,26 | 4,0858 | 4 | |

| | | | | | | | | | | | | | | |
|---|---|---|---|---|---|---|---|---|---|---|---|---|---|---|
| Q15149-4;Q15149- | 3192;3160; | Q15149-4 | Q15149-4 | Plectin | PLEC | DAKAYSDPSTGEP | >sp\|Q15149-4\|PLEC_ | 3037 | 1 | 2,2106 | 3,4156 | 2,47 | 3,6834 | 4 |
| A0A0C4DG51;Q9NF | 573;573;61 | A0A0C4DG | A0A0C4DG51 | Calcium-independe | PNPLA8 | LGTGRYESDVRNT | >tr\|A0A0C4DG51\|A0 | 1025 | 1 | 4,4073 | 5,1474 | 4,18 | 5,2803 | 4 |
| P30041 | 89 | P30041 | P30041 | Peroxiredoxin-6 | PRDX6 | DSVEDHLAWSKD | >sp\|P30041\|PRDX6_ | 2555 | 1 | 4,361 | 5,0396 | 5,1 | 4,9938 | 4 |
| A6NLN1;P26599;P2 | 97;127;127 | A6NLN1 | A6NLN1 | Polypyrimidine trac | PTBP1 | NYYTSVTPVLRGQ | >tr\|A6NLN1\|A6NLN1 | 1203 | 1 | 4,4968 | 4,8022 | 2,16 | 4,0766 | 4 |
| P18031 | 46 | P18031 | P18031 | Tyrosine-protein ph | PTPN1 | PCRVAKLPKNKNF | >sp\|P18031\|PTN1_H | 2442 | 1 | 4,9152 | 2,9615 | 3,46 | 2,2989 | 4 |
| H0YC15;Q05209 | 27;64 | H0YC15 | H0YC15 | Tyrosine-protein ph | PTPN12 | PTATGEKEENVKK | >tr\|H0YC15\|H0YC15_ | 1778 | 1 | 4,1774 | 5,166 | 2,8 | 3,0043 | 4 |
| A8MXQ1;P53801 | 144;165 | A8MXQ1 | A8MXQ1 | Pituitary tumor-trar | PTTG1IP | RRAEMKTRHDEIF | >tr\|A8MXQ1\|A8MXQ | 1230 | 1;2 | 6,4829 | 6,056 | 5,52 | 6,3435 | 4 |
| A8MXQ1;P53801;A | 153;174;10 | A8MXQ1 | A8MXQ1 | Pituitary tumor-trar | PTTG1IP | DEIRKKYGLFKEEN | >tr\|A8MXQ1\|A8MXQ | 1229 | 1;2 | 3,0552 | 2,4328 | 2,45 | 2,7111 | 4 |
| O14966 | 54 | O14966 | O14966 | Ras-related protein | RAB7L1 | TVGVDFALKVLQV | >sp\|O14966\|RAB7L_ | 2052 | 1 | 3,7529 | 6,3043 | 3,38 | 6,2782 | 4 |
| P54136-2;P54136 | 312;384 | P54136-2 | P54136-2 | Arginine--tRNA ligas | RARS | CSIPLTIVKSDGGY | >sp\|P54136-2\|SYRC_ | 2703 | 1 | 2,8921 | 4,2034 | 3,49 | 4,1627 | 4 |
| H7C3J0;Q6NUM9 | 275;486 | H7C3J0 | H7C3J0 | All-trans-retinol 13, | RETSAT | FEEWQAELKGKR | >tr\|H7C3J0\|H7C3J0_ | 1876 | 1 | 3,1801 | 2,6208 | 3,41 | 2,3327 | 4 |
| J3QR17;J3KSI3;C9JE | 34;34;34;3 | J3QR17 | J3QR17 | E3 ubiquitin-protei | RFFL | PPPQGARMQAYS | >tr\|J3QR17\|J3QR17_ | 1380 | 1 | 4,4631 | 5,3965 | 2,15 | 5,5595 | 4 |
| H0YE35;Q9NVN3-4; | 18;202;402 | H0YE35 | H0YE35 | Synembryn-B;Syner | RIC8A;RIC8B | FLFVLCSESVPRFI | >tr\|H0YE35\|H0YE35_ | 1314 | 1 | 4,7237 | 4,4926 | 4,8 | 4,7773 | 4 |
| P39023;B5MCW2;G | 307;255;25 | P39023 | P39023 | 60S ribosomal prot | RPL3 | LIKDGKLIKNNAST | >sp\|P39023\|RL3_HU | 2604 | 1 | 2,7011 | 3,198 | 3,65 | 3,5417 | 4 |
| C9J9K3;P08865;A0 | 139;139;14 | C9J9K3 | C9J9K3 | 40S ribosomal prot | RPSA | VTDPRADHQPLTE | >tr\|C9J9K3\|C9J9K3_ | 1022 | 1 | 4,3856 | 4,5825 | 2,7 | 2,3454 | 4 |
| Q8N122-3;Q8N122 | 565;723 | Q8N122-3 | Q8N122-3 | Regulatory-associat | RPTOR | VRDSPCTPRLRSV | >sp\|Q8N122-3\|RPTO | 3222 | 1 | 2,8057 | 8,136 | 3,69 | 8,3349 | 4 |
| V9GYG0;P12236;P1 | 112;112;11 | V9GYG0 | V9GYG0 | ADP/ATP transloca | SLC25A6;SLC25A4 | QLFLGGVDRHKQ | >tr\|V9GYG0\|V9GYG0 | 2360 | 1 | 2,8523 | 4,6344 | 3,06 | 4,2393 | 4 |
| O95295 | 129 | O95295 | O95295 | SNARE-associated | SNAPIN | AKETARRRAMLD | >sp\|O95295\|SNAPN_ | 2173 | 1;2 | 4,325 | 4,7676 | 4,74 | 5,4672 | 4 |
| P00966;Q5T6L6 | 133;133 | P00966 | P00966 | Argininosuccinate s | ASS1 | ATGKGNDQVRFE | >sp\|P00966\|ASSY_HU | 2207 | 1 | 4,0195 | 3,6721 | 3,84 | 3,3784 | 4 |
| Q9UBP0-3;Q9UBP0 | 126;212 | Q9UBP0-3 | Q9UBP0-3 | Spastin | SPAST | KMQPVLPFSKSQ | >sp\|Q9UBP0-3\|SPAS | 3591 | 1 | 2,8764 | 3,5844 | 3,45 | 4,1694 | 4 |
| P12931;P12931-2 | 530;536 | P12931 | P12931 | Proto-oncogene tyr | SRC | LQAFLEDYFTSTEF | >sp\|P12931\|SRC_HU | 2369 | 1 | 5,2448 | 4,8519 | 6,93 | 5,5991 | 4 |
| P12931;P12931-2 | 329;335 | P12931 | P12931 | Proto-oncogene tyr | SRC | AQVMKKLRHEKL | >sp\|P12931\|SRC_HU | 2372 | 1 | 3,3424 | 3,9429 | 3,94 | 4,2617 | 4 |
| P12931;P12931-2 | 187;193 | P12931 | P12931 | Proto-oncogene tyr | SRC | RGTFLVRESETTK | >sp\|P12931\|SRC_HU | 2368 | 1 | 5,7722 | 8,9221 | 5,61 | 8,4694 | 4 |
| P12931;P12931-2 | 93;93 | P12931 | P12931 | Proto-oncogene tyr | SRC | RAGPLAGGVTTF | >sp\|P12931\|SRC_HU | 2367 | 1 | 7,1512 | 6,7369 | 6,09 | 7,0363 | 4 |
| P12931;P12931-2 | 216;222 | P12931 | P12931 | Proto-oncogene tyr | SRC | NVKHYKIRKLDSG | >sp\|P12931\|SRC_HU | 2370 | 1 | 7,2096 | 6,3528 | 4,41 | 5,9677 | 4 |
| Q14849-2;Q14849- | 254;272;27 | Q14849-2 | Q14849-2 | StAR-related lipid tr | STARD3 | ENWKFEKNNEYG | >sp\|Q14849-2\|STAR3 | 1957 | 1 | 5,0568 | 8,1851 | 4,14 | 7,3569 | 4 |
| J3KQA0;P21579 | 227;230 | J3KQA0 | J3KQA0 | Synaptotagmin-1 | SYT1 | VPYSELGGKTLVM | >tr\|J3KQA0\|J3KQA0_ | 1920 | 1 | 2,7476 | 2,7434 | 2,71 | 3,4311 | 4 |
| Q96C24 | 400 | Q96C24 | Q96C24 | Synaptotagmin-like | SYTL4 | YADEAKKRSNPYV | >sp\|Q96C24\|SYTL4_ | 3351 | 1 | 5,288 | 5,6603 | 6,05 | 5,9894 | 4 |
| Q5T011-5;Q5T011 | 2070;2127 | Q5T011-5 | Q5T011-5 | Protein SZT2 | SZT2 | ALPPSLALSRSQEF | >sp\|Q5T011-5\|SZT2_ | 3110 | 1 | 4,777 | 5,2793 | 4,03 | 4,116 | 4 |
| Q7L7X3 | 610 | Q7L7X3 | Q7L7X3 | Serine/threonine-p | TAOK1 | FQAEEEANLLRRQ | >sp\|Q7L7X3\|TAOK1_ | 3177 | 1 | 3,0164 | 2,3286 | 3,26 | 2,8964 | 4 |
| G3V133;Q66K14-2; | 29;853;853 | G3V133 | G3V133 | TBC1 domain family | TBC1D9B | GCSRTMAGRRDP | >tr\|G3V133\|G3V133_ | 1714 | 1 | 5,3323 | 4,0529 | 4,08 | 3,0199 | 4 |

| | | | | | | | | | | | | | | | |
|---|---|---|---|---|---|---|---|---|---|---|---|---|---|---|---|
| A0A087WXC6;E9PF | 79;138;174 | A0A087WX | A0A087WXC6 | Transducin beta-lik | TBL2 | DTLRVFKMTKRED | >tr\|A0A087WXC6\|A0 | 901 | | 1 | 3,8874 | 3,7857 | 4,14 | 3,8586 | 4 |
| P37173;P37173-2 | 424;449 | P37173 | P37173 | TGF-beta receptor t | TGFBR2 | SVDDLANSGQVG | >sp\|P37173\|TGFR2_H | 2598 | | 1 | 4,0176 | 3,8387 | 6,44 | 3,1597 | 4 |
| P21980-2;P21980 | 369;369 | P21980-2 | P21980-2 | Protein-glutamine g | TGM2 | WQALDPTPQEKS | >sp\|P21980-2\|TGM2_ | 2473 | | 1 | 4,9858 | 5,4983 | 5,37 | 5,9493 | 4 |
| C9JZ87;Q9NUM4 | 84;84 | C9JZ87 | C9JZ87 | Transmembrane pr | TMEM106B | RIPRGQENQLVAL | >tr\|C9JZ87\|C9JZ87_H | 1408 | | 1 | 5,4103 | 6,94 | 4,85 | 6,5032 | 4 |
| C9JZ87;Q9NUM4;F2 | 18;18;18 | C9JZ87 | C9JZ87 | Transmembrane pr | TMEM106B | KSLSHLPLHSSKED | >tr\|C9JZ87\|C9JZ87_H | 1407 | | 1 | 4,668 | 6,5697 | 4,9 | 6,5382 | 4 |
| C9JZ87;Q9NUM4;F2 | 50;50;50 | C9JZ87 | C9JZ87 | Transmembrane pr | TMEM106B | VHNEDGRNGDVS | >tr\|C9JZ87\|C9JZ87_H | 1409 | | 1 | 3,972 | 5,3802 | 3,47 | 4,9765 | 4 |
| F8W120;F8VWX5;F | 40;40;40;4 | F8W120 | F8W120 | Transmembrane pr | TMEM106C | LLAEREQEEAIAQ | >tr\|F8W120\|F8W120 | 917 | | 1 | 4,1066 | 4,1156 | 3,56 | 3,8613 | 4 |
| Q8IY95-2;Q8IY95 | 126;130 | Q8IY95-2 | Q8IY95-2 | Transmembrane pr | TMEM192 | QYHHSKIRNRGYN | >sp\|Q8IY95-2\|TM192 | 3216 | | 1 | 3,4323 | 5,2441 | 3,85 | 5,5923 | 4 |
| Q8IY95-2;Q8IY95 | 240;244 | Q8IY95-2 | Q8IY95-2 | Transmembrane pr | TMEM192 | SSLEEIVEKQGDTI | >sp\|Q8IY95-2\|TM192 | 3217 | | 1 | 4,2796 | 6,083 | 3,91 | 6,4794 | 4 |
| Q8IY95-2;Q8IY95 | 209;213 | Q8IY95-2 | Q8IY95-2 | Transmembrane pr | TMEM192 | NKAKPEPDILEEEK | >sp\|Q8IY95-2\|TM192 | 3213 | 1;2 | | 4,3009 | 5,1802 | 2,97 | 3,924 | 4 |
| Q8IY95-2;Q8IY95 | 211;215 | Q8IY95-2 | Q8IY95-2 | Transmembrane pr | TMEM192 | AKPEPDILEEEKIYA | >sp\|Q8IY95-2\|TM192 | 3214 | 1;2 | | 7,289 | 6,3585 | 5,82 | 5,0244 | 4 |
| Q86T03;Q86T03-2 | 146;153 | Q86T03 | Q86T03 | Transmembrane pr | TMEM55B | LICKVTSQRIACPR | >sp\|Q86T03\|TM55B_ | 3185 | | 1 | 7,6538 | 9,1669 | 8,29 | 8,8989 | 4 |
| E9PCX8;Q68CZ2;Q6 | 958;855;61 | E9PCX8 | E9PCX8 | Tensin-3 | TNS3 | SMCSTPAFPVSPE | >tr\|E9PCX8\|E9PCX8_ | 1552 | 1;2 | | 4,1573 | 2,0921 | 4,29 | 3,5204 | 4 |
| Q15785 | 54 | Q15785 | Q15785 | Mitochondrial impo | TOMM34 | LQAQGSSDPEEES | >sp\|Q15785\|TOM34_ | 3077 | | 1 | 2,3564 | 4,3631 | 2,74 | 4,4025 | 4 |
| O95801 | 129 | O95801 | O95801 | Tetratricopeptide r | TTC4 | DLNAVLYTNRAAA | >sp\|O95801\|TTC4_H | 2186 | | 1 | 3,3931 | 2,3275 | 2,25 | 3,3766 | 4 |
| P68363;P68363-2;P | 282;166;26 | P68363 | P68363 | Tubulin alpha-1B ch | TUBA1B;TUBA4A | FPLATYAPVISAEK | >sp\|P68363\|TBA1B_H | 2812 | | 1 | 8,3434 | 5,7152 | 6,98 | 4,5244 | 4 |
| F2Z2B9;Q9BSJ2;Q9 | 83;83;83 | F2Z2B9 | F2Z2B9 | Gamma-tubulin cor | TUBGCP2 | DELKSKNTRNLDP | >tr\|F2Z2B9\|F2Z2B9_H | 1631 | | 1 | 3,6219 | 3,1715 | 3,64 | 2,4488 | 4 |
| E9PKD3;E2QRB9;Q1 | 131;93;93; | E9PKD3 | E9PKD3 | Thioredoxin reduct | TXNRD1 | YRVALREKKVVYE | >tr\|E9PKD3\|E9PKD3_ | 825 | | 1 | 5,2425 | 3,5307 | 2,74 | 2,9233 | 4 |
| Q9P2Y5-2;E9PR71;Q | 52;323;424 | Q9P2Y5-2 | Q9P2Y5-2 | UV radiation resista | UVRAG | EFPLYPKGGEKLQ | >sp\|Q9P2Y5-2\|UVRA | 1623 | | 1 | 3,0357 | 5,6299 | 3,29 | 5,7214 | 4 |
| P51809-3;P51809;P | 49;90;90 | P51809-3 | P51809-3 | Vesicle-associated r | VAMP7 | AFNFLNEIKKRFQT | >sp\|P51809-3\|VAMP | 2687 | | 1 | 3,1144 | 4,6231 | 3,88 | 4,8368 | 4 |
| B4DWR3;P61758 | 175;180 | B4DWR3 | B4DWR3 | Prefoldin subunit 3 | VBP1 | LRDQFTTTEVNMA | >tr\|B4DWR3\|B4DWR | 1303 | | 1 | 3,2521 | 3,0035 | 4,03 | 2,7575 | 4 |
| Q9H270;A0A087WX | 4;4 | Q9H270 | Q9H270 | Vacuolar protein so | VPS11 | _____________M | >sp\|Q9H270\|VPS11_ | 910 | | 1 | 2,6965 | 2,4462 | 2,95 | 2,0648 | 4 |
| S4R3Q6;O75436-2; | 132;157;15 | S4R3Q6 | S4R3Q6 | Vacuolar protein so | VPS26A | DLVKEYDLIVHQLA | >tr\|S4R3Q6\|S4R3Q6_ | 2140 | | 1 | 7,5539 | 4,2393 | 6,5 | 2,7393 | 4 |
| Q96QK1 | 507 | Q96QK1 | Q96QK1 | Vacuolar protein so | VPS35 | GRFIHLLRSEDPD | >sp\|Q96QK1\|VPS35_ | 3392 | | 1 | 4,9961 | 3,6479 | 5,19 | 4,0491 | 4 |
| F5H4M0;Q9H9H4 | 189;191 | F5H4M0 | F5H4M0 | Vacuolar protein so | VPS37B | LPPRLPELAPTAPL | >tr\|F5H4M0\|F5H4M0 | 1657 | | 1 | 3,9666 | 2,8971 | 3,89 | 2,5432 | 4 |
| Q9UEU0-2;Q9UEU0 | 7;68;82 | Q9UEU0-2 | Q9UEU0-2 | Vesicle transport th | VTI1B | __________MEEE | >sp\|Q9UEU0-2\|VTI1B | 3593 | | 1 | 5,907 | 2,8011 | 6,21 | 2,1753 | 4 |
| H3BSK9;Q8WWM7- | 289;349;34 | H3BSK9 | H3BSK9 | Ataxin-2-like protei | ATXN2L | GSGRESPSLASRE | >tr\|H3BSK9\|H3BSK9_ | 1832 | | 1 | 5,7713 | 3,3531 | 6,86 | 3,5973 | 4 |
| O75083;O75083-3 | 238;98 | O75083 | O75083 | WD repeat-containi | WDR1 | KVCALGGSKAHD | >sp\|O75083\|WDR1_H | 2129 | | 1 | 4,1796 | 5,5339 | 2,82 | 4,9595 | 4 |
| Q6PJI9-3;Q6PJI9;Q6 | 425;833;27 | Q6PJI9-3 | Q6PJI9-3 | WD repeat-containi | WDR59 | WGESSPEELRFGS | >sp\|Q6PJI9-3\|WDR59 | 3156 | | 1 | 3,0937 | 5,7132 | 2,1 | 5,4587 | 4 |
| S4R3J8;Q9Y5A9 | 37;37 | S4R3J8 | S4R3J8 | YTH domain family | YTHDF2 | SVHQKDGLNDDD | >tr\|S4R3J8\|S4R3J8_H | 3670 | | 1 | 5,7479 | 2,9311 | 5,79 | 3,1282 | 4 |
| Q68DK2-2;Q68DK2; | 1875;1896; | Q68DK2-2 | Q68DK2-2 | Zinc finger FYVE do | ZFYVE26 | EKPEALDSSKSESP | >sp\|Q68DK2-2\|ZFY26 | 3140 | | 1 | 3,5824 | 3,6586 | 3,99 | 3,4678 | 4 |

| | | | | | | | | | | | | | | |
|---|---|---|---|---|---|---|---|---|---|---|---|---|---|---|
| P56945-5;P56945-8 | 222;224;22 | P56945-5 | P56945-5 | Breast cancer anti-e | BCAR1 | PAKVVVPTRVGQ | >sp\|P56945-5\|BCAR1 | 2725 | 1;2 | 2,5271 | 3,374 | 3,15 | 2,9718 | 4 |
| K7ELY9;E7EV84;Q1 | 107;157;23 | K7ELY9 | K7ELY9 | Beclin-1 | BECN1 | EAERLDQEEAQYC | >tr\|K7ELY9\|K7ELY9_ | 1526 | 1 | 2,0507 | 3,1174 | 2,6 | 3,3894 | 4 |
| F5GXW6;O95415 | 57;57 | F5GXW6 | F5GXW6 | Brain protein I3 | BRI3 | YPYLVTGIPTHHPI | >tr\|F5GXW6\|F5GXW | 1642 | 1 | 2,9395 | 3,4676 | 3,51 | 3,6076 | 4 |
| Q8WUW1;Q8WUW | 63;63 | Q8WUW1 | Q8WUW1 | Protein BRICK1 | BRK1 | ATLNEKLTALERRI | >sp\|Q8WUW1\|BRK1_ | 3281 | 1 | 2,981 | 4,2833 | 3,3 | 5,0349 | 4 |
| Q15067-3;Q15067- | 591;629;62 | Q15067-3 | Q15067-3 | Peroxisomal acyl-co | ACOX1 | DVTLGSVLGRYDG | >sp\|Q15067-3\|ACOX | 3033 | 1 | 5,0133 | 7,5846 | 5,81 | 7,5967 | 4 |
| P49589-3;E9PLP0;P | 115;115;32 | P49589-3 | P49589-3 | Cysteine--tRNA liga | CARS | QWSPPAGTQPCR | >sp\|P49589-3\|SYCC_ | 2651 | 1 | 4,7456 | 2,9128 | 2,77 | 2,8612 | 4 |
| P24666-3;P24666-2 | 98;132;132 | P24666-3 | P24666-3 | Low molecular weig | ACP1 | LGSYDPQKQLIIED | >sp\|P24666-3\|PPAC_ | 2507 | 1 | 6,3395 | 2,0442 | 5,76 | 2,5108 | 4 |
| D6R9B4;Q04900-4; | 160;175;18 | D6R9B4 | D6R9B4 | Sialomucin core pro | CD164 | VIFFLYKFCKSKERI | >tr\|D6R9B4\|D6R9B4_ | 1456 | 1 | 3,8771 | 2,2974 | 3,54 | 2,2513 | 4 |
| E5RIH6;E5RJT5;E5R | 78;78;78;7 | E5RIH6 | E5RIH6 | Centrosomal protei | CEP170 | LNGTFVNDVRIPE | >tr\|E5RIH6\|E5RIH6_ | 1481 | 1 | 3,1245 | 3,9666 | 3,81 | 4,2162 | 4 |
| Q53EZ4;H0Y432;Q5 | 237;77;237 | Q53EZ4 | Q53EZ4 | Centrosomal protei | CEP55 | KPESEGYLQEEKQ | >sp\|Q53EZ4\|CEP55_ | 3089 | 1 | 2,9318 | 3,1163 | 3,78 | 2,1971 | 4 |
| Q00610-2;Q00610; | 899;899;90 | Q00610-2 | Q00610-2 | Clathrin heavy chai | CLTC | IDSNNNPERFLREI | >sp\|Q00610-2\|CLH1_ | 876 | 1 | 2,4687 | 5,2008 | 3,27 | 5,5074 | 4 |
| P60709;P63261;I3L | 218;218;20 | P60709;P6 | P60709 | Actin, cytoplasmic 1 | ACTB;ACTG1;POTE | TAEREIVRDIKEKL | >sp\|P60709\|ACTB_H | 2735 | 1 | 5,3561 | 5,2055 | 4,86 | 5,1583 | 4 |
| G5E9W3;Q9UKF6 | 640;677 | G5E9W3 | G5E9W3 | Cleavage and polya | CPSF3 | DESLREMVELAAC | >tr\|G5E9W3\|G5E9W | 1738 | 1 | 4,5985 | 2,8305 | 2,06 | 2,8174 | 4 |
| P60709;G5E9R0;E7 | 91;91;91;9 | P60709;P6 | P60709 | Actin, cytoplasmic 1 | ACTB;ACTG1;POTE | ITNWDDMEKIWH | >sp\|P60709\|ACTB_H | 2734 | 1 | 6,0446 | 4,5157 | 3,92 | 3,0163 | 4 |
| Q9UJU6-5;H0Y5J4; | 121;153;22 | Q9UJU6-5 | Q9UJU6-5 | Drebrin-like protein | DBNL | RRERELREAARRE | >sp\|Q9UJU6-5\|DBNL | 1766 | 1 | 2,5588 | 2,101 | 3,98 | 2,7008 | 4 |
| G3V4I3;F8VNZ9;G3 | 22;22;22;2 | G3V4I3 | G3V4I3 | Diacylglycerol kinas | DGKA | LISPSDFAQLQKYN | >tr\|G3V4I3\|G3V4I3_ | 1392 | 1 | 3,436 | 2,47 | 4,01 | 2,8067 | 4 |
| Q12959-8;Q12959- | 283;283;34 | Q12959-8 | Q12959-8 | Disks large homolog | DLG1 | HEEAVTALKNTSD | >sp\|Q12959-8\|DLG1_ | 1013 | 1 | 3,5497 | 5,9404 | 3,83 | 6,1558 | 4 |
| Q08495-3;Q08495- | 268;253;29 | Q08495-3 | Q08495-3 | Dematin | DMTN | TSLHQGTSKSSSLI | >sp\|Q08495-3\|DEMA | 2890 | 1 | 2,5557 | 4,8023 | 2,76 | 5,2452 | 4 |
| P50570-3;P50570-2 | 593;593;59 | P50570-3 | P50570-3 | Dynamin-2 | DNM2 | NKHVFAIFNTEQR | >sp\|P50570-3\|DYN2_ | 2676 | 1 | 3,098 | 3,0143 | 4,28 | 3,2258 | 4 |
| Q14126 | 979 | Q14126 | Q14126 | Desmoglein-2 | DSG2 | TERVYAPASTLVD | >sp\|Q14126\|DSG2_H | 2974 | 1;2 | 5,6677 | 2,7761 | 6,39 | 3,0503 | 4 |
| F8WAW4;F1LLU7;C | 81;111;158 | F8WAW4 | F8WAW4 | Enoyl-CoA delta iso | ECI2 | WNALGSLPKEAAI | >tr\|F8WAW4\|F8WAV | 1027 | 1 | 2,9209 | 3,1433 | 3,43 | 2,7809 | 4 |
| F8WAW4;F1LLU7;C | 36;66;113; | F8WAW4 | F8WAW4 | Enoyl-CoA delta iso | ECI2 | KLLKKDPGNEVKL | >tr\|F8WAW4\|F8WAV | 1026 | 1 | 5,9409 | 3,2972 | 3,52 | 2,0975 | 4 |
| Q9BSW2;Q9BSW2- | 141;141 | Q9BSW2 | Q9BSW2 | EF-hand calcium-bi | EFCAB4B | EDAGEQVAQRHE | >sp\|Q9BSW2\|EFC4B_ | 3448 | 1 | 4,1312 | 2,5935 | 4,67 | 2,9595 | 4 |
| Q99613-2;Q99613; | 871;881;88 | Q99613-2 | Q99613-2 | Eukaryotic translati | EIF3C;EIF3CL | LVENNERVFDHK | >sp\|Q99613-2\|EIF3C_ | 1312 | 1 | 2,4911 | 2,8761 | 3,11 | 3,2907 | 4 |
| P06733;K7EM90;P1 | 44;44;44;4 | P06733 | P06733 | Alpha-enolase;Enol | ENO1;ENO3;ENO2 | GLFRAAVPSGAST | >sp\|P06733\|ENOA_H | 2254 | 1 | 2,1996 | 2,6971 | 2,02 | 2,947 | 4 |
| Q96TA1-2;Q96TA1 | 384;397 | Q96TA1-2 | Q96TA1-2 | Niban-like protein 1 | FAM129B | IDKLGEYMEKLSRI | >sp\|Q96TA1-2\|NIBL1 | 3404 | 1 | 2,578 | 2,4526 | 3,37 | 2,1294 | 4 |
| I3L2L5;C9JLW8;I3L | 36;41;41;6 | I3L2L5 | I3L2L5 | Protein FAM195B | FAM195B | SEIFTPAHEENVRF | >tr\|I3L2L5\|I3L2L5_HU | 1391 | 1 | 4,5395 | 3,61 | 3,91 | 3,5343 | 4 |
| A0A0A0MTJ1;Q143 | 205;205;36 | A0A0A0M | A0A0A0MTJ1 | Peptidyl-prolyl cis-t | FKBP8 | LVKKHAAQRSTET | >tr\|A0A0A0MTJ1\|A0 | 994 | 1 | 3,1201 | 2,6867 | 4,81 | 3,5312 | 4 |
| A0A140T905;A2ABJ | 65;128;153 | A0A140T9 | A0A140T905 | Flotillin-1 | FLOT1 | EMAKAQRDYELKI | >tr\|A0A140T905\|A0A | 1143 | 1 | 2,4288 | 2,8629 | 3,18 | 3,2453 | 4 |
| P06241-3;P06241-2 | 385;437;44 | P06241-3; | P12931 | Tyrosine-protein kir | FYN;YES1;SRC | GAKFPIKWTAPEA | >sp\|P06241-3\|FYN_H | 2253 | 1 | 3,5099 | 3,997 | 3,82 | 3,8715 | 4 |
| P11413;P11413-3;P | 503;533;54 | P11413 | P11413 | Glucose-6-phospha | G6PD | GPTEADELMKRV | >sp\|P11413\|G6PD_H | 2352 | 1 | 4,9232 | 2,3617 | 3,81 | 2,3683 | 4 |

| | | | | | | | | | | | | | | |
|---|---|---|---|---|---|---|---|---|---|---|---|---|---|---|
| Q9NXN4-2;Q9NXN4 | 236;236 | Q9NXN4-2 | Q9NXN4-2 | Ganglioside-induce | GDAP2 | LYFPRSLKEENRSL | >sp\|Q9NXN4-2\|GDAP | 3563 | 1 | 6,2596 | 6,8045 | 6,69 | 7,3808 | 4 |
| Q5TBH8;O15228 | 10;10 | Q5TBH8 | Q5TBH8 | Dihydroxyacetone p | GNPAT | ______MESSSSSI | >tr\|Q5TBH8\|Q5TBH8 | 2061 | 1 | 3,3065 | 8,5487 | 4,27 | 8,2826 | 4 |
| P17174-2;P17174 | 243;264 | P17174-2 | P17174-2 | Aspartate aminotra | GOT1 | FEFFCAQSFSKNF | >sp\|P17174-2\|AATC_ | 2437 | 1 | 2,3482 | 3,73 | 2,04 | 3,6559 | 4 |
| F5H0Q1;F5H234;F5 | 207;213;20 | F5H0Q1 | F5H0Q1 | Integral membrane | GPR137 | ACLCLVARRAPST | >tr\|F5H0Q1\|F5H0Q1_ | 1640 | 1 | 4,3041 | 2,8729 | 2,77 | 3,2215 | 4 |
| Q8NFJ5 | 347 | Q8NFJ5 | Q8NFJ5 | Retinoic acid-induc | GPRC5A | QKEFSIPRAHAWF | >sp\|Q8NFJ5\|RAI3_HU | 3255 | 1;2 | 5,3727 | 7,3057 | 7,68 | 5,2339 | 4 |
| Q8NFJ5 | 300 | Q8NFJ5 | Q8NFJ5 | Retinoic acid-induc | GPRC5A | KPQLVKKSYGVEN | >sp\|Q8NFJ5\|RAI3_HU | 3257 | 1;2;3 | 3,1309 | 6,1388 | 3,02 | 5,8163 | 4 |
| Q8NFJ5 | 350 | Q8NFJ5 | Q8NFJ5 | Retinoic acid-induc | GPRC5A | FSIPRAHAWPSPY | >sp\|Q8NFJ5\|RAI3_HU | 3256 | 1;2 | 5,8827 | 6,2085 | 4,73 | 5,0259 | 4 |
| Q8NFJ5 | 320 | Q8NFJ5 | Q8NFJ5 | Retinoic acid-induc | GPRC5A | ITQGFEETGDTLY | >sp\|Q8NFJ5\|RAI3_HU | 3259 | 1;2 | 3,6219 | 5,7463 | 2,42 | 5,2709 | 4 |
| Q8NFJ5 | 317 | Q8NFJ5 | Q8NFJ5 | Retinoic acid-induc | GPRC5A | QEEITQGFEETGD | >sp\|Q8NFJ5\|RAI3_HU | 3258 | 1;2 | 3,6769 | 4,2171 | 1,66 | 3,9851 | 4 |
| Q9NQ84;Q9NQ84-2 | 438;450;48 | Q9NQ84 | Q9NQ84 | G-protein coupled r | GPRC5C | TPPKDGKNSQVFF | >sp\|Q9NQ84\|GPC5C_ | 1016 | 1 | 5,5608 | 5,3163 | 4,79 | 8,2463 | 4 |
| Q9NQ84;Q9NQ84-2 | 324;336;36 | Q9NQ84 | Q9NQ84 | G-protein coupled r | GPRC5C | EQSYQGDMYPTR | >sp\|Q9NQ84\|GPC5C_ | 1015 | 1 | 4,5646 | 3,6523 | 4,72 | 3,5948 | 4 |
| Q9NQ84;Q9NQ84-2 | 358;370;40 | Q9NQ84 | Q9NQ84 | G-protein coupled r | GPRC5C | FSMDEPVAAKRP | >sp\|Q9NQ84\|GPC5C_ | 1017 | 1;2 | 5,8441 | 8,3063 | 5,16 | 1,499 | 4 |
| P48637-2;P48637 | 361;472 | P48637-2 | P48637-2 | Glutathione synthe | GSS | ADGGVAAGVAVL | >sp\|P48637-2\|GSHB_ | 2643 | 1 | 3,608 | 3,0795 | 3,36 | 2,8673 | 4 |
| P53701 | 51 | P53701 | P53701 | Cytochrome c-type | HCCS | PVNTEPSGPTCEK | >sp\|P53701\|CCHL_HU | 2701 | 1 | 5,141 | 3,0196 | 6,84 | 3,356 | 4 |
| P53701 | 61 | P53701 | P53701 | Cytochrome c-type | HCCS | CEKKTYSVPAHQE | >sp\|P53701\|CCHL_HU | 2699 | 1 | 2,9518 | 2,8075 | 3,75 | 3,4725 | 4 |
| P19367-4;P19367-2 | 720;731;73 | P19367-4 | P19367-4 | Hexokinase-1 | HK1 | CLDDIRTHYDRLVI | >sp\|P19367-4\|HXK1_ | 2462 | 1 | 2,7903 | 2,2868 | 3,49 | 3,8492 | 4 |
| P04075;J3KPS3;H3E | 3;3;3;57;3; | P04075 | P04075 | Fructose-bisphosph | ALDOA | ______________M | >sp\|P04075\|ALDOA_ | 1914 | 1 | 2,2783 | 2,4125 | 2,86 | 2,4414 | 4 |
| G3V159;Q9UPZ3-2; | 105;105;21 | G3V159 | G3V159 | Hermansky-Pudlak | HPS5 | REKFWKIGNKERD | >tr\|G3V159\|G3V159_ | 1716 | 1 | 5,9338 | 5,0512 | 6,31 | 5,5132 | 4 |
| Q86YV9 | 410 | Q86YV9 | Q86YV9 | Hermansky-Pudlak | HPS6 | ELPSAKDLVFEEA | >sp\|Q86YV9\|HPS6_H | 3202 | 1 | 4,4555 | 5,2208 | 4,51 | 5,5567 | 4 |
| Q92835-2;Q92835; | 914;915;70 | Q92835-2 | Q92835-2 | Phosphatidylinosito | INPP5D | PPCSGSSITEIINPN | >sp\|Q92835-2\|SHIP1 | 3312 | 1 | 4,4065 | 3,2347 | 1,25 | 2,8339 | 3 |
| O15357;O15357-2; | 986;744;92 | O15357 | O15357 | Phosphatidylinosito | INPPL1 | GVAAPPPKNSFNI | >sp\|O15357\|SHIP2_H | 2068 | 1;2 | 2,2029 | 2,4365 | 2,31 | 1,9777 | 3 |
| Q15058 | 1023 | Q15058 | Q15058 | Kinesin-like protein | KIF14 | IEELEKAKQHLEQE | >sp\|Q15058\|KIF14_H | 3031 | 1 | 2,4014 | 2,5407 | 1,03 | 2,1544 | 3 |
| Q96L93-5;Q96L93-6 | 923;1079;1 | Q96L93-5 | Q96L93-5 | Kinesin-like protein | KIF16B | QERLEYEIQQLKQ | >sp\|Q96L93-5\|KI16B_ | 3373 | 1 | 3,0957 | 1,9911 | 3,25 | 2,1763 | 3 |
| Q6IAA8 | 140 | Q6IAA8 | Q6IAA8 | Ragulator complex | LAMTOR1 | PFSDLQQVSRIAA | >sp\|Q6IAA8\|LTOR1_H | 3144 | 1;2 | 6,9398 | 3,9326 | 6,4 | 1,994 | 3 |
| Q6IAA8;H0YFI1;F5H | 40;27;40;4 | Q6IAA8 | Q6IAA8 | Ragulator complex | LAMTOR1 | PSSPPTKALNGAE | >sp\|Q6IAA8\|LTOR1_H | 3142 | 1 | 4,1251 | 2,238 | 1,67 | 2,1833 | 3 |
| C9JXK9;C9JUT4;A0A | 138;301;27 | C9JXK9 | C9JXK9 | Lipoma-preferred p | LPP | GYGYAPNQGRYY | >tr\|C9JXK9\|C9JXK9_H | 929 | 1 | 5,5217 | 1,2488 | 5,06 | 3,5505 | 3 |
| Q96AG4 | 203 | Q96AG4 | Q96AG4 | Leucine-rich repeat | LRRC59 | RKREKAEEKERRR | >sp\|Q96AG4\|LRC59_ | 3343 | 1 | 1,8699 | 3,5264 | 2,36 | 4,0778 | 3 |
| O95297-4;O95297-1 | 117;240;24 | O95297-4 | O95297-4 | Myelin protein zero | MPZL1 | LVKSLPSGSHQGP | >sp\|O95297-4\|MPZL1 | 2176 | 1 | 2,3156 | 6,9746 | 1,41 | 4,9573 | 3 |
| J3KRL6;O94916-2;C | 44;220;296 | J3KRL6 | J3KRL6 | Nuclear factor of ac | NFAT5 | ELKIVVQPETQHR | >tr\|J3KRL6\|J3KRL6_H | 1923 | 1 | 1,6934 | 4,5437 | 2,87 | 4,2679 | 3 |
| Q96CV9-3;Q96CV9- | 299;350;35 | Q96CV9-3 | Q96CV9-3 | Optineurin | OPTN | NSAIPSELNEKQEI | >sp\|Q96CV9-3\|OPTN | 3354 | 1 | 2,145 | 3,1746 | 1,93 | 3,1264 | 3 |
| O00264 | 113 | O00264 | O00264 | Membrane-associat | PGRMC1 | FDVTKGRKFYGPE | >sp\|O00264\|PGRC1_ | 2018 | 1 | 2,5835 | 1,3441 | 2,89 | 2,0378 | 3 |

| | | | | | | | | | | | | | | | |
|---|---|---|---|---|---|---|---|---|---|---|---|---|---|---|---|
| Q9NRX4 | 125 | Q9NRX4 | Q9NRX4 | 14 kDa phosphohist | PHPT1 | KAKYPDYEVTWAI | >sp\|Q9NRX4\|PHP14_ | 3544 | | 1 | 1,5879 | 2,5261 | 2,84 | 3,3111 | 3 |
| P42356 | 1154 | P42356 | P42356 | Phosphatidylinositc | PI4KA | FMASLNLRNRYA( | >sp\|P42356\|PI4KA_H | 2618 | | 1 | 1,2068 | 2,6883 | 2,88 | 3,7177 | 3 |
| Q15149-4;Q15149- | 4380;4348; | Q15149-4 | Q15149-4 | Plectin | PLEC | KTKMSAAQALKK( | >sp\|Q15149-4\|PLEC_ | 3041 | | 1 | 2,5443 | 3,3403 | 1,88 | 3,4071 | 3 |
| H0Y8J0;Q08752 | 16;323 | H0Y8J0 | H0Y8J0 | Peptidyl-prolyl cis-t | PPID | KALYRRAQGWQ( | >tr\|H0Y8J0\|H0Y8J0_ | 1770 | | 1 | 2,4474 | 3,3523 | 2,01 | 1,7622 | 3 |
| Q05655;Q05655-2 | 313;313 | Q05655 | Q05655 | Protein kinase C de | PRKCD | ASRRSDSASSEPV( | >sp\|Q05655\|KPCD_H | 2868 | 1;2 | | 4,6789 | 2,4976 | 3,76 | 1,9553 | 3 |
| E5RFX7;E5RG77;O9 | 17;69;69 | E5RFX7 | E5RFX7 | Proline synthase co | PROSC | VIEAYGHGQRTFG | >tr\|E5RFX7\|E5RFX7_ | 1482 | | 1 | 3,0501 | 1,7474 | 3,46 | 2,0508 | 3 |
| Q6P2Q9 | 2091 | Q6P2Q9 | Q6P2Q9 | Pre-mRNA-processi | PRPF8 | RAISAANLHLRTN | >sp\|Q6P2Q9\|PRP8_H | 3151 | | 1 | 3,4448 | 5,9871 | 1,91 | 6,0147 | 3 |
| Q9Y520-3;Q9Y520- | 985;985;98 | Q9Y520-3 | Q9Y520-3 | Protein PRRC2C | PRRC2C | EGFIRSSEGPKPEK | >sp\|Q9Y520-3\|PRC2( | 1505 | | 1 | 1,0041 | 3,6588 | 2,03 | 2,6989 | 3 |
| O00232-2;O00232 | 117;137 | O00232-2 | O00232-2 | 26S proteasome no | PSMD12 | LRLIDTLRMVTEGI | >sp\|O00232-2\|PSD12 | 2016 | | 1 | 1,8996 | 5,234 | 2,59 | 3,3944 | 3 |
| Q05397-7;Q05397; | 815;861;82 | Q05397-7 | Q05397-7 | Focal adhesion kina | PTK2 | REDGSLQGPIGN( | >sp\|Q05397-7\|FAK1_ | 2857 | | 1 | 2,8595 | 7,1848 | 1,8 | 4,6143 | 3 |
| P11216 | 76 | P11216 | P11216 | Glycogen phosphor | PYGB | RDHLVGRWIRTQ | >sp\|P11216\|PYGB_H | 2345 | | 1 | 1,6172 | 2,6357 | 2,29 | 3,2807 | 3 |
| A0A087WZ85;A0A( | 815;812;89 | A0A087WZ | A0A087WZ85 | Roundabout homol | ROBO1 | WLYRHRKKRNGL | >tr\|A0A087WZ85\|A0 | 835 | | 1 | 4,0603 | 1,8258 | 4,12 | 2,4552 | 3 |
| O15027-2;O15027- | 813;813;81 | O15027-2 | O15027-2 | Protein transport p | SEC16A | PDGNKANHSSHQ | >sp\|O15027-2\|SC16A | 1628 | | 1 | 3,0731 | 5,9078 | 1,84 | 5,4727 | 3 |
| P61619-3;P61619;B | 156;276;28 | P61619-3 | P61619-3 | Protein transport p | SEC61A1 | FRVDLPIKSARYR( | >sp\|P61619-3\|S61A1 | 1296 | | 1 | 3,2351 | 2,6045 | 2,99 | 1,4417 | 3 |
| Q15427 | 16 | Q15427 | Q15427 | Splicing factor 3B su | SF3B4 | MAAGPISERNQD | >sp\|Q15427\|SF3B4_ | 3065 | | 1 | 1,5531 | 2,1058 | 3,06 | 3,1689 | 3 |
| Q9BRG2-2;Q9BRG2 | 109;231 | Q9BRG2-2 | Q9BRG2-2 | SH2 domain-contai | SH2D3A | TPSFELPDASERPI | >sp\|Q9BRG2-2\|SH23 | 3442 | | 2 | 3,1931 | 3,7274 | 1,04 | 3,5287 | 3 |
| P34897-3;P34897;P | 271;292;28 | P34897-3 | P34897-3 | Serine hydroxymetl | SHMT2 | TTHKTLRGARSGL | >sp\|P34897-3\|GLYM_ | 2581 | | 1 | 1,4962 | 3,0981 | 1,77 | 3,6378 | 3 |
| B6ZDF2;H7BYB7;Q8 | 126;212;26 | B6ZDF2 | B6ZDF2 | Solute carrier famil | SLC15A4 | PPDGSAFTDMFKI | >tr\|B6ZDF2\|B6ZDF2_ | 1313 | | 1 | 2,0768 | 2,2314 | 1,89 | 2,2574 | 3 |
| O60264 | 413 | O60264 | O60264 | SWI/SNF-related m | SMARCA5 | ADVEKSLPPKKEV | >sp\|O60264\|SMCA5_ | 2107 | | 1 | 3,5525 | 3,0309 | 4,38 | 1,8831 | 3 |
| Q07617 | 660 | Q07617 | Q07617 | Sperm-associated a | SPAG1 | KYSECLKINNKEC/ | >sp\|Q07617\|SPAG1_ | 2880 | | 1 | 1,81 | 3,4876 | 2,92 | 4,1542 | 3 |
| O60271-5;O60271- | 138;138;13 | O60271-5 | O60271-5 | C-Jun-amino-termir | SPAG9;MAPK8IP3 | SLESQTRQLELKAI | >sp\|O60271-5\|JIP4_ | 958 | | 1 | 1,689 | 4,5839 | 3,08 | 4,619 | 3 |
| A6NMU3;Q92783-2 | 108;219;21 | A6NMU3 | A6NMU3 | Signal transducing a | STAM | LLTNHQHEGRKVI | >tr\|A6NMU3\|A6NMU | 1206 | | 1 | 1,2651 | 3,5501 | 3,03 | 3,1177 | 3 |
| Q14849-2;Q14849- | 279;297;29 | Q14849-2 | Q14849-2 | StAR-related lipid tr | STARD3 | TFILKTFLPCPAEL\ | >sp\|Q14849-2\|STAR3 | 1958 | | 1 | 3,0417 | 2,7947 | 1,16 | 2,8111 | 3 |
| E9PNS3;F2Z2Y8;Q9 | 13;13;13 | E9PNS3 | E9PNS3 | Toll-interacting pro | TOLLIP | ___MATTVSTQR( | >tr\|E9PNS3\|E9PNS3_ | 1614 | | 1 | 2,6354 | 2,0888 | 3,11 | 1,4086 | 3 |
| P22314-2;P22314;( | 20;60;60;7 | P22314-2 | P22314-2 | Ubiquitin-like modi | UBA1 | GSEADIDEGLYSR( | >sp\|P22314-2\|UBA1_ | 2477 | | 1 | 1,3244 | 2,0043 | 3,76 | 3,1025 | 3 |
| P68036-2;P68036;P | 115;147;20 | P68036-2 | P68036-2 | Ubiquitin-conjugati | UBE2L3 | DRKKFCKNAEEFT | >sp\|P68036-2\|UB2L3 | 2804 | | 1 | 2,4981 | 1,176 | 4 | 2,3318 | 3 |
| Q9UHR4 | 163 | Q9UHR4 | Q9UHR4 | Brain-specific angio | BAIAP2L1 | QGSRNALKYEHKE | >sp\|Q9UHR4\|BI2L1_ | 3603 | | 1 | 1,1189 | 3,6281 | 2,09 | 3,7875 | 3 |
| G8JLQ3;P78537-2;P | 66;116;144 | G8JLQ3 | G8JLQ3 | Biogenesis of lysosc | BLOC1S1 | IELDMRTIATALEY | >tr\|G8JLQ3\|G8JLQ3_ | 1744 | | 1 | 2,1151 | 1,9301 | 2,69 | 3,4256 | 3 |
| P17655 | 66 | P17655 | P17655 | Calpain-2 catalytic s | CAPN2 | FPAIPSALGFKELG | >sp\|P17655\|CAN2_H | 2438 | | 1 | 1,7327 | 3,7955 | 2,22 | 3,692 | 3 |
| A0A0U1RR39;A0A1 | 630;672;67 | A0A0U1RR | A0A0U1RR39 | E3 ubiquitin-protei | CBL | CDHPKIKPSSSAN/ | >tr\|A0A0U1RR39\|A0/ | 1138 | | 1 | 2,0662 | 2,2144 | 2,05 | 1,0547 | 3 |
| P50990-2;P50990;I | 11;30;22 | P50990-2 | P50990-2 | T-complex protein : | CCT8 | _____MHFSGLEE | >sp\|P50990-2\|TCPQ_ | 2677 | | 1 | 5,1946 | 2,3966 | 2,5 | 1,6252 | 3 |

| | | | | | | | | | | | | | | | |
|---|---|---|---|---|---|---|---|---|---|---|---|---|---|---|---|
| P24666-3;P24666-2 | 109;143;14 | P24666-3 | P24666-3 | Low molecular weig | ACP1 | IEDPYYGNDSDFE | >sp\|P24666-3\|PPAC_ | 2509 | | 1 | 2,1878 | 4,5008 | 1,1 | 3,7087 | 3 |
| Q9ULV4;Q9ULV4-2; | 301;307;35 | Q9ULV4 | Q9ULV4 | Coronin-1C;Coronin | CORO1C | GDSSIRYFEITDESI | >sp\|Q9ULV4\|COR1C_ | 3617 | | 1 | 3,8956 | 3,4235 | 1,61 | 2,9224 | 3 |
| P41240;H3BN15;H3 | 18;18;18 | P41240 | P41240 | Tyrosine-protein kin | CSK | AIQAAWPSGTECI | >sp\|P41240\|CSK_HU | 2611 | | 1 | 5,9662 | 2,906 | 4,56 | 1,2857 | 3 |
| P12814-2;P12814;P | 193;193;19 | P12814-2 | P12814-2 | Alpha-actinin-1 | ACTN1 | GFCALIHRHRPELI | >sp\|P12814-2\|ACTN1 | 2364 | | 1 | 2,8671 | 2,7084 | 2,05 | 1,6993 | 3 |
| P07108;P07108-3;B | 29;30;39;4 | P07108 | P07108 | Acyl-CoA-binding pr | DBI | RHLKTKPSDEEML | >sp\|P07108\|ACBP_H | 993 | | 1 | 4,9938 | 4,0851 | 3,12 | 1,591 | 3 |
| X6RA56;X6RCK5;O7 | 72;65;72;7 | X6RA56 | X6RA56 | Dynactin subunit 3 | DCTN3 | LYKKIEDLIKYLDPE | >tr\|X6RA56\|X6RA56_ | 978 | | 1 | 1,829 | 3,0553 | 2,41 | 3,3171 | 3 |
| H3BPN2;H3BSA6;Q | 8;30;129 | H3BPN2 | H3BPN2 | dCTP pyrophosphat | DCTPP1 | ________MDINR | >tr\|H3BPN2\|H3BPN2 | 1826 | | 1 | 1,4745 | 2,5542 | 2,36 | 2,4399 | 3 |
| P15924;P15924-2;P | 1065;1065; | P15924 | P15924 | Desmoplakin | DSP | ENCNKNKFLDQN | >sp\|P15924\|DESP_HU | 2423 | | 1 | 3,0764 | 2,0291 | 3,21 | 1,9204 | 3 |
| M0QZW4;M0R248; | 149;146;14 | M0QZW4 | M0QZW4 | Delta(3,5)-Delta(2,4 | ECH1 | DVARISWYLRDIIT | >tr\|M0QZW4\|M0QZ | 2003 | | | 1,7066 | 2,848 | 2,04 | 2,139 | 3 |
| X6RAC9;A6NJH9;P4 | 78;89;106; | X6RAC9 | X6RAC9 | Eukaryotic translati | EIF1AY;EIF1AX | VILKYNADEARSLK | >tr\|X6RAC9\|X6RAC9_ | 1201 | | 1 | 2,5725 | 2,1383 | 2,18 | 1,364 | 3 |
| Q5JVZ5;Q96JJ3;Q5J | 717;719;28 | Q5JVZ5 | Q5JVZ5 | Engulfment and cel | ELMO2 | PPIPKEPSSYDFVY | >tr\|Q5JVZ5\|Q5JVZ5_ | 3103 | | 1 | 3,6996 | 5,9965 | 1,5 | 6,4238 | 3 |
| H7BXK9;Q9NP58-4; | 314;420;46 | H7BXK9 | H7BXK9 | ATP-binding cassett | ABCB6 | ARAVDSLLNFETV | >tr\|H7BXK9\|H7BXK9_ | 1853 | | 1 | 1,7834 | 2,2213 | 2,57 | 2,3154 | 3 |
| A0A087WWY3;Q5H | 2282;2574; | A0A087WW | A0A087WWY3 | Filamin-A | FLNA | TPCEEILVKHVGSF | >tr\|A0A087WWY3\|A | 897 | | 1 | 3,3671 | 3,6867 | 1,91 | 3,9538 | 3 |
| Q13155 | 35 | Q13155 | Q13155 | Aminoacyl tRNA syn | AIMP2 | LPTCMYRLPNVH | >sp\|Q13155\|AIMP2_ | 2934 | | 1 | 2,3552 | 6,2856 | 1,11 | 6,6902 | 3 |
| Q8NFJ5;H0YFN2 | 293;20 | Q8NFJ5 | Q8NFJ5 | Retinoic acid-induce | GPRC5A | PVEDAFCKPQLVK | >sp\|Q8NFJ5\|RAI3_HU | 3260 | | 1 | 5,8668 | 6,9407 | 5,52 | 6,1671 | 3 |
| Q9NQ84;Q9NQ84-2 | 387;399;43 | Q9NQ84 | Q9NQ84 | G-protein coupled r | GPRC5C | PTEMALMHKVPS | >sp\|Q9NQ84\|GPC5C_ | 1021 | 1;2 | | 3,365 | 3,269 | 3,42 | 3,1626 | 3 |
| P00390-5;P00390-4 | 353;382;39 | P00390-5 | P00390-5 | Glutathione reducta | GSR | PIGTVGLTEDEAIH | >sp\|P00390-5\|GSHR_ | 2195 | | 1 | 1,7645 | 2,3922 | 3,17 | 2,1225 | 3 |
| Q53GQ0-2;A0A1B0 | 66;34;66;6 | Q53GQ0-2 | Q53GQ0-2 | Estradiol 17-beta-de | HSD17B12 | EWAVVTGSTDGI | >sp\|Q53GQ0-2\|DHB1 | 1176 | | 1 | 1,1786 | 3,6853 | 2,34 | 3,9958 | 3 |
| G5E9S2;E7ET17;E7E | 593;596;70 | G5E9S2 | G5E9S2 | Peroxisomal multifu | HSD17B4 | GNIMLSQKLQMII | >tr\|G5E9S2\|G5E9S2_ | 1519 | | 1 | 1,5023 | 2,2076 | 2,28 | 2,2655 | 3 |
| P11142;E9PKE3;E9P | 525;506;37 | P11142 | P11142 | Heat shock cognate | HSPA8 | LSKEDIERMVQEA | >sp\|P11142\|HSP7C_H | 2338 | | 1 | 1,5861 | 2,8045 | 2,45 | 2,0234 | w |
| P26639;P26639-2 | 284;317 | P26639 | P26639 | Threonine--tRNA lig | TARS | TGKIKALKIHKNSS | >sp\|P26639\|SYTC_HU | 2524 | | 1 | 1,5993 | 1,8414 | 2,28 | 2,9084 | w |
| K7EJH3;A0A1B0GW | 4;4;4;4;4;4 | K7EJH3 | K7EJH3 | Splicing factor U2AF | U2AF1L4;U2AF1 | _____________M | >tr\|K7EJH3\|K7EJH3_H | 1179 | | 1 | 1,2974 | 1,7915 | 2,24 | 2,9483 | w |
| P38646 | 196 | P38646 | P38646 | Stress-70 protein, m | HSPA9 | YLGHTAKNAVITV | >sp\|P38646\|GRP75_H | 2600 | | 1 | 1,9362 | 1,8469 | 2,28 | 1,0666 | u |
| H7BYN4;Q02241 | 736;736 | H7BYN4 | H7BYN4 | Kinesin-like protein | KIF23 | SVASCISEWEQKIF | >tr\|H7BYN4\|H7BYN4 | 1858 | | 1 | 1,0897 | 2,3968 | 1,78 | 1,7189 | u |
| P00338;P00338-4;P | 239;181;26 | P00338 | P00338 | L-lactate dehydroge | LDHA | KEQWKEVHKQV | >sp\|P00338\|LDHA_H | 2192 | | 1 | 2,6268 | 2,4676 | 3,02 | 2,6118 | u |
| P07355;P07355-2;H | 235;253;23 | P07355 | P07355 | Annexin A2;Annexin | ANXA2;ANXA2P2 | RSVPHLQKVFDRY | >sp\|P07355\|ANXA2_ | 2271 | | 1 | 1,82 | 1,332 | 1,95 | 2,2834 | u |
| Q15149-4;Q15149- | 3640;3608; | Q15149-4 | Q15149-4 | Plectin | PLEC | IIEKTEIIRQQGLAS | >sp\|Q15149-4\|PLEC_ | 3047 | | 1 | 1,0756 | 1,1691 | 2,17 | 1,0059 | u |
| E9PNN3;Q03393 | 65;133 | E9PNN3 | E9PNN3 | 6-pyruvoyl tetrahyd | PTS | LQKVLPVGVLYKV | >tr\|E9PNN3\|E9PNN3 | 1613 | | 1 | 2,5401 | 1,1234 | 1,06 | 1,3408 | u |
| S4R3Q3;P20336 | 28;123 | S4R3Q3 | S4R3Q3 | Ras-related protein | RAB3A | ESFNAVQDWSTQ | >tr\|S4R3Q3\|S4R3Q3_ | 2467 | | 1 | 1,1976 | 1,3272 | 1,84 | 2,7165 | u |
| P61006-2;P61006 | 143;143 | P61006-2 | P61006-2 | Ras-related protein | RAB8A | KRQVSKERGEKLA | >sp\|P61006-2\|RAB8A | 2740 | | 1 | 1,2149 | 1,1376 | 2,03 | 1,1716 | u |
| O43567-2;O43567; | 256;375;17 | O43567-2 | O43567-2 | E3 ubiquitin-protein | RNF13 | EHDVVVQLQPNG | >sp\|O43567-2\|RNF13 | 2096 | | 1 | 1,6892 | 1,3792 | 2,39 | 1,1209 | u |

| | | | | | | | | | | | | | | |
|---|---|---|---|---|---|---|---|---|---|---|---|---|---|---|
| P49591;Q5T5C7 | 410;410 | P49591 | P49591 | Serine--tRNA ligase, | SARS | CSNCTDYQARRLF | >sp\|P49591\|SYSC_HU | 2653 | 1 | 1,7398 | 1,2475 | 2,67 | 1,1466 | u |
| Q9BWJ5 | 18 | Q9BWJ5 | Q9BWJ5 | Splicing factor 3B su | SF3B5 | DRYTIHSQLEHLQ! | >sp\|Q9BWJ5\|SF3B5_ | 3455 | 1 | 1,5214 | 2,2422 | 1,71 | 1,1851 | u |
| P05141 | 165 | P05141 | P05141 | ADP/ATP transloca: | SLC25A5 | AEREFRGLGDCLV | >sp\|P05141\|ADT2_H | 2233 | | 1,0406 | 1,0699 | 2,13 | 1,3165 | u |
| Q13813;Q13813-2;. | 2423;2428; | Q13813 | Q13813 | Spectrin alpha chai | SPTAN1 | EIESAFRALSSEGK | >sp\|Q13813\|SPTN1_I | 1084 | 1 | 1,5043 | 2,4931 | 1,09 | 1,6551 | u |
| Q13242;H0YIB4;S4I | 70;58;70 | Q13242 | Q13242 | Serine/arginine-rich | SRSF9 | AFVRFEDPRDAED | >sp\|Q13242\|SRSF9_I | 2940 | 1 | 1,37 | 1,3186 | 2,39 | 1,1718 | u |
| P29401;P29401-2;A | 275;283;10 | P29401 | P29401 | Transketolase | TKT | KPLPKNMAEQIIQ | >sp\|P29401\|TKT_HUI | 2554 | 1 | 1,7161 | 2,1055 | 1,31 | 1,5606 | u |
| O75674-3;J3KQU4;I | 32;109;102 | O75674-3 | O75674-3 | TOM1-like protein : | TOM1L1 | KEFVKENLVKLLNI | >sp\|O75674-3\|TM1L: | 1896 | 1 | 1,9881 | 1,8141 | 2,79 | 1,7343 | u |
| H0YMH8;H0YKJ9;O | 35;154;179 | H0YMH8 | H0YMH8 | Tetraspanin-3 | TSPAN3 | LCACIVLCRRSRDI | >tr\|H0YMH8\|H0YMH | 1802 | 1;2 | 2,1137 | 1,5523 | 1,98 | 1,9023 | u |
| Q9C0H2-3;Q9C0H2- | 338;477;50 | Q9C0H2-3 | Q9C0H2-3 | Protein tweety hon | TTYH3 | RESPPPSYTSSMR. | >sp\|Q9C0H2-3\|TTYH: | 3474 | 1 | 1,2294 | 1,4717 | 2,29 | 1,3708 | u |
| A6NJA2;P54578-2;F | 4;4;4;4 | A6NJA2 | A6NJA2 | Ubiquitin carboxyl-I | USP14 | ____________M | >tr\|A6NJA2\|A6NJA2_ | 1199 | 1 | 2,844 | 1,3365 | 1,33 | 1,1837 | u |
| P26640-2;A0A140T! | 133;347;42 | P26640-2 | P26640-2 | Valine--tRNA ligase | VARS | LQEVWKWKEEKC | >sp\|P26640-2\|SYVC_ | 1144 | 1 | 1,6258 | 1,6907 | 2,03 | 1,4375 | u |
| Q9UEU0-2;Q9UEUC | 54;115;32;: | Q9UEU0-2 | Q9UEU0-2 | Vesicle transport th | VTI1B | LTATPGGRGDMK | >sp\|Q9UEU0-2\|VTI1E | 3595 | 1 | 3,0075 | 1,7194 | 1,22 | 1,8132 | u |
| Q68DK2-2;Q68DK2; | 1712;1733; | Q68DK2-2 | Q68DK2-2 | Zinc finger FYVE dor | ZFYVE26 | DSLLSRYAEKALDI | >sp\|Q68DK2-2\|ZFY26 | 3139 | 1 | 1,3542 | 1,1863 | 2,48 | 1,7132 | u |
| O00154-2;K7EKP8;( | 152;135;15 | O00154-2 | O00154-2 | Cytosolic acyl coen: | ACOT7 | VVYSRQEQEEEGI | >sp\|O00154-2\|BACH_ | 1973 | 1 | 1,3387 | 1,651 | 2,79 | 1,7815 | u |
| P16152 | 194 | P16152 | P16152 | Carbonyl reductase | CBR1 | TKKGVHQKEGWI | >sp\|P16152\|CBR1_HI | 2431 | 1 | 1,2493 | 1,6389 | 3,6 | 1,8617 | u |
| Q13045-2;Q13045-: | 1120;1164; | Q13045-2 | Q13045-2 | Protein flightless-1 | FLII | YMKHTRLFRCSNI | >sp\|Q13045-2\|FLII_H | 2929 | 1 | 1,2146 | 1,5209 | 2,45 | 1,2815 | u |
| P09211;A8MX94;A( | 64;64;52 | P09211 | P09211 | Glutathione S-trans | GSTP1 | LYGQLPKFQDGDI | >sp\|P09211\|GSTP1_I | 2322 | 1 | 1,2747 | 1,4179 | 1,37 | 2,9026 | u |
| P12081-4;P12081;E | 65;65;65;6! | P12081-4 | P12081-4 | Histidine--tRNA liga | HARS | SKQKFVLKTPKGT | >sp\|P12081-4\|SYHC_ | 2356 | 1 | 1,461 | 1,4043 | 1,48 | 2,1809 | u |
| E9PC71;P84074;P3: | 58;58;58 | E9PC71 | E9PC71 | Neuron-specific cal | HPCAL1;HPCA | LTVDEFKKIYANFF | >tr\|E9PC71\|E9PC71_ | 1550 | 1 | 1,8393 | 5,4137 | 1,3 | 6,0231 | u |
| P07355;P07355-2;I | 188;206;18 | P07355 | P07355 | Annexin A2;Annexir | ANXA2 | ALAKGRRAEDGS\ | >sp\|P07355\|ANXA2_ | 2263 | 1 | 1,3506 | 5,7435 | 1,28 | 5,4551 | R2 |
| H0YHW7;Q7Z4F1-2 | 376;474;47 | H0YHW7 | H0YHW7 | Low-density lipopr | LRP10 | ALGCTCKLYAIRTC | >tr\|H0YHW7\|H0YHW | 1797 | 1 | 1,8006 | 2,584 | 1,03 | 2,8245 | R2 |
| Q8N565;Q8N565-2 | 37;37 | Q8N565 | Q8N565 | Melanoregulin | MREG | RALPEKEPLVSDN | >sp\|Q8N565\|MREG_ | 3236 | 1 | 1,2052 | 4,2472 | 1,37 | 2,7601 | R2 |
| Q01968-2;Q01968 | 477;477 | Q01968-2 | Q01968-2 | Inositol polyphosph | OCRL | AFVDFNEGEIKFIP | >sp\|Q01968-2\|OCRL_ | 2843 | 1 | 1,8493 | 3,5311 | 1,68 | 4,9504 | R2 |
| Q13131;Q13131-2 | 442;457 | Q13131 | Q13131 | 5'-AMP-activated p | PRKAA1 | AIKQLDYEWKVVI | >sp\|Q13131\|AAPK1_ | 2931 | 1 | 1,0313 | 2,3014 | 1,4 | 3,2896 | R2 |
| Q15262;Q15262-2;( | 831;832;83 | Q15262 | Q15262 | Receptor-type tyro: | PTPRK | PLSITFMDQHNFS | >sp\|Q15262\|PTPRK_I | 3056 | 1 | 1,5137 | 5,2219 | 1,71 | 5,6241 | R2 |
| P06737-2;E9PK47;P | 130;164;16 | P06737-2 | P06737-2 | Glycogen phosphor | PYGL | ATLGLAAYGYGIR\ | >sp\|P06737-2\|PYGL_ | 1593 | 1 | 1,5187 | 3,1609 | 1,86 | 2,9124 | R2 |
| Q9P258 | 359 | Q9P258 | Q9P258 | Protein RCC2 | RCC2 | VLDSQKRVFSWG | >sp\|Q9P258\|RCC2_H | 3584 | 1 | 1,8965 | 2,2186 | 1,84 | 2,4294 | R2 |
| Q9HB90 | 5 | Q9HB90 | Q9HB90 | Ras-related GTP-bir | RRAGC | ___________MS | >sp\|Q9HB90\|RRAGC_ | 3520 | 1 | 1,9176 | 3,7621 | 1,44 | 2,4768 | R2 |
| P34897-3;P34897;P | 207;228;21 | P34897-3 | P34897-3 | Serine hydroxymetl | SHMT2 | ARLFRPRLIIAGTS. | >sp\|P34897-3\|GLYM_ | 2580 | 1 | 1,5878 | 5,3627 | 2,72 | 4,7401 | R2 |
| Q9Y2Z0-2;Q9Y2Z0 | 245;277 | Q9Y2Z0-2 | Q9Y2Z0-2 | Suppressor of G2 al | SUGT1 | DVPTPKQFVADVI | >sp\|Q9Y2Z0-2\|SGT1_ | 3639 | 1 | 1,1716 | 3,0572 | 1,34 | 3,3578 | R2 |
| H0Y547;P98196;E9I | 243;268;26 | H0Y547 | H0Y547 | Probable phospholi | ATP11A | GATLKNTEKIFGV. | >tr\|H0Y547\|H0Y547_ | 1568 | 1 | 1,7014 | 3,2891 | 1,58 | 5,0509 | R2 |

| | | | | | | | | | | | | | | |
|---|---|---|---|---|---|---|---|---|---|---|---|---|---|---|
| J3KRD5;A0A087WV | 83;87;126;: | J3KRD5 | J3KRD5 | Cytosolic non-speci | CNDP2 | WDSEPFTLVERD( | >tr\|J3KRD5\|J3KRD5_I | 881 | 1 | 1,4428 | 4,0048 | 1,78 | 3,9001 | R2 |
| P17844-2;P17844;J: | 123;202;20 | P17844-2 | P17844-2 | Probable ATP-depe | DDX5;DDX17 | AAEYCRACRLKST( | >sp\|P17844-2\|DDX5_ | 1940 | 1 | 1,0905 | 2,6926 | 1,66 | 5,1749 | R2 |
| P55039;J3QLF3;J3Q | 37;37;37;3 | P55039 | P55039 | Developmentally-re | DRG2 | ATEYHLGLLKAKL/ | >sp\|P55039\|DRG2_H | 2713 | 1 | 1,9194 | 5,8847 | 1,98 | 6,2886 | R2 |
| P50402;Q5HY57 | 90;55 | P50402 | P50402 | Emerin | EMD | DLPKKEDALLYQSI | >sp\|P50402\|EMD_HU | 2668 | 1 | 1,6293 | 3,4379 | 1,35 | 3,077 | R2 |
| P11413;P11413-3;P | 507;537;55 | P11413 | P11413 | Glucose-6-phospha | G6PD | ADELMKRVGFQY | >sp\|P11413\|G6PD_H | 2353 | 1 | 1,3358 | 5,8661 | 1,97 | 6,7473 | R2 |
| A0A087WXK4;Q7Z3 | 777;805 | A0A087WX | A0A087WXK4 | Integral membrane | GPR155 | LIEVGLASDRGEA\ | >tr\|A0A087WXK4\|A0 | 908 | 1 | 1,4154 | 2,4945 | 1,31 | 2,0381 | R2 |
| Q92835-2;Q92835 | 220;221 | Q92835-2 | Q92835-2 | Phosphatidylinositc | INPP5D | LPHLKKLTTLLCKE | >sp\|Q92835-2\|SHIP1_ | 3316 | 1 | 2,1931 | 1,9189 | 2,97 | 1,9553 | R1 |
| P26006 | 1028 | P26006 | P26006 | Integrin alpha-3;Int | ITGA3 | LWKCGFFKRARTI | >sp\|P26006\|ITA3_HU | 2519 | 1 | 2,004 | 1,3714 | 2,12 | 1,2041 | R1 |
| J3QRK0;P16144-4;P | 16;1399;13 | J3QRK0;P1 | P16144-4 | Integrin beta-4 | ITGB4 | STNSLHRMTTTSA | >tr\|J3QRK0\|J3QRK0_ | 1961 | 1;2 | 5,9327 | 1,9924 | 4,71 | 1,6426 | R1 |
| P02545-6;P02545-3 | 45;45;45;4! | P02545-6 | P02545-6 | Prelamin-A/C;Lamir | LMNA | QEKEDLQELNDRL | >sp\|P02545-6\|LMNA_ | 2215 | 1 | 4,3538 | 1,6679 | 5,85 | 1,4038 | R1 |
| O75197;A4QPB2-2; | 518;136;13 | O75197 | O75197 | Low-density lipopro | LRP5;LRP5L | GWPNGLALDLQE | >sp\|O75197\|LRP5_HI | 2136 | 1 | 3,0417 | 1,4026 | 2,79 | 1,5073 | R1 |
| F8VRL4;F8VRR0;A0, | 16;16;16;1( | F8VRL4 | F8VRL4 | Diphosphoinositol p | NUDT4;NUDT11;N | MEPEEEPGGAAV | >tr\|F8VRL4\|F8VRL4_I | 1032 | 1 | 3,668 | 1,3725 | 3,48 | 1,3963 | R1 |
| Q96BI3-3;Q96BI3 | 186;256 | Q96BI3-3 | Q96BI3-3 | Gamma-secretase s | APH1A | RSLLCRRQEDSRV | >sp\|Q96BI3-3\|APH1A | 3347 | 1 | 2,2801 | 1,038 | 3,01 | 1,6609 | R1 |
| Q14155-1;B1ALK7;( | 595;654;68 | Q14155-1 | Q14155-1 | | ARHGEF7 | NGQTVIEEKSLVD | >sp\|Q14155-1\|ARHG | 1259 | 1 | 4,7474 | 1,8752 | 4,5 | 1,9591 | R1 |
| Q9H2G2-2;Q9H2G2 | 1194;1225 | Q9H2G2-2 | Q9H2G2-2 | STE20-like serine/tl | SLK | SECLNPSTQSRISK | >sp\|Q9H2G2-2\|SLK_I | 3496 | 1 | 3,948 | 1,2119 | 4,16 | 1,1379 | R1 |
| Q9NTJ3-2;E9PD53;( | 616;591;61 | Q9NTJ3-2 | Q9NTJ3-2 | Structural maintena | SMC4 | DAIIQEKKSGRIPG | >sp\|Q9NTJ3-2\|SMC4_ | 1559 | 1 | 2,1475 | 1,2997 | 3,04 | 1,3591 | R1 |
| P12931;P12931-2 | 379;385 | P12931 | P12931 | Proto-oncogene tyr | SRC | PQLVDMAAQIAS( | >sp\|P12931\|SRC_HU | 2371 | 1 | 4,1229 | 1,6241 | 3,52 | 1,5963 | R1 |
| Q96A25-2;B7Z779;I | 34;82;82;8: | Q96A25-2 | Q96A25-2 | Transmembrane pr | TMEM106A | KIPQELEKQLVALI | >sp\|Q96A25-2\|T106/ | 1325 | 1 | 4,3216 | 1,9469 | 4,03 | 1,7203 | R1 |
| Q8IZQ1-2;Q8IZQ1 | 1944;1944 | Q8IZQ1-2 | Q8IZQ1-2 | WD repeat and FYV | WDFY3 | KAFAADTGMNRS | >sp\|Q8IZQ1-2\|WDFY: | 3219 | 1 | 2,5415 | 2,08 | 3,52 | 1,1062 | R1 |
| O60716-13;O60716 | 226;226;22 | O60716-13 | O60716-13 | Catenin delta-1 | CTNND1 | VGGSSVDLHRFHI | >sp\|O60716-13\|CTNI | 1414 | 1 | 3,3354 | 1,3296 | 2,42 | 1,5786 | R1 |
| Q8TDM6-2;Q8TDM | 429;319;42 | Q8TDM6-2 | Q8TDM6-2 | Disks large homolo | DLG5 | TAKESEKYREERD/ | >sp\|Q8TDM6-2\|DLG5 | 3270 | 1 | 4,1255 | 1,2974 | 4,76 | 1,5492 | R1 |
| P49588;P49588-2;I | 667;675;11 | P49588 | P49588 | Alanine--tRNA ligas | AARS | AEEIANEMIEAAK. | >sp\|P49588\|SYAC_HU | 2649 | 1 | 5,0998 | 1,3827 | 6,24 | 1,1355 | R1 |
| Q9Y262-2;Q9Y262;I | 491;539;58 | Q9Y262-2 | Q9Y262-2 | Eukaryotic translati | EIF3L | DKDMIHIADTKV/ | >sp\|Q9Y262-2\|EIF3L_ | 1237 | 1 | 2,7787 | 1,1276 | 2,33 | 1,2023 | R1 |
| Q8TBA6-2;Q8TBA6 | 36;36 | Q8TBA6-2 | Q8TBA6-2 | Golgin subfamily A | GOLGA5 | GAATALSRKDNAS | >sp\|Q8TBA6-2\|GOGA | 3264 | 1 | 2,8955 | 1,4325 | 3,85 | 1,6794 | R1 |
| P09211;A8MX94;A( | 50;50;38 | P09211 | P09211 | Glutathione S-trans | GSTP1 | TVETWQEGSLKAS | >sp\|P09211\|GSTP1_I | 2321 | | 2,0782 | 1,2324 | 4 | 1,8349 | R1 |
| G3V547;G3V4D9;G: | 30;30;30;3( | G3V547 | G3V547 | Nuclear export mec | NEMF | AELNASLLGMRVI | >tr\|G3V547\|G3V547_ | 1723 | 1 | 1,7272 | 1,3055 | 1,43 | 1,5262 | E |
| P19174;P19174-2 | 771;771 | P19174 | P19174 | 1-phosphatidylinosi | PLCG1 | INEEALEKIGTAEP | >sp\|P19174\|PLCG1_I | 2456 | 1 | 1,1031 | 1,4988 | 1,42 | 1,943 | E |
| P26639;P26639-2 | 298;331 | P26639 | P26639 | Threonine--tRNA lig | TARS | TYWEGKADMETL | >sp\|P26639\|SYTC_HU | 2523 | 1 | 1,5051 | 1,7096 | 1,8 | 1,5804 | E |
| Q9P2Y5-2;E9PR71;( | 55;326;427 | Q9P2Y5-2 | Q9P2Y5-2 | UV radiation resista | UVRAG | LYPKGGEKLQFDY | >sp\|Q9P2Y5-2\|UVRA( | 1624 | 1 | 1,9446 | 1,4877 | 1,05 | 1,9518 | E |
| P46108;P46108-2 | 136;136 | P46108 | P46108 | Adapter molecule c | CRK | SRQGSGVILRQEE | >sp\|P46108\|CRK_HU | 2634 | 1 | 1,2167 | 1,2249 | 1,58 | 1,1713 | E |
| Q96C19 | 83 | Q96C19 | Q96C19 | EF-hand domain-co | EFHD2 | GIGEPQSPSRRVFI | >sp\|Q96C19\|EFHD2_ | 3350 | 1 | 1,2356 | 1,7587 | 1,65 | 1,3861 | E |

| | | | | | | | | | | | | | | | |
|---|---|---|---|---|---|---|---|---|---|---|---|---|---|---|---|
| A0FGR8-5;H7BXI1;A | 231;787;79 | A0FGR8-5 | A0FGR8-5 | Extended synaptota | ESYT2 | ACRNLIAFSEDGSI | >sp\|A0FGR8-5\|ESYT2 | 911 | | 1 | 1,7093 | 1,2362 | 1,76 | 1,3652 | E |
| H0YF56;E9PPK2;Q6 | 5;153;251;2 | H0YF56 | H0YF56 | Phosphofurin acidic | PACS1 | ___________UIK | >tr\|H0YF56\|H0YF56_ | 1618 | | 1 | 3,9503 | 3,9414 | 1,75 | 1,6512 | A |
| P00966;Q5T6L6;Q5 | 87;87;87 | P00966 | P00966 | Argininosuccinate s | ASS1 | FIWPAIQSSALYED | >sp\|P00966\|ASSY_HU | 2209 | | 1 | 2,1363 | 2,6639 | 1,22 | 1,5349 | A |
| O60784-4;O60784;O | 108;141;14 | O60784-4 | O60784-4 | Target of Myb prote | TOM1;UNQ1844 | DAFRSSPDLTGVV | >sp\|O60784-4\|TOM1 | 2120 | | 1 | 2,0956 | 5,8076 | 1,21 | 1,5467 | A |
| H7C086;P48047;H7 | 35;35;35 | H7C086 | H7C086 | ATP synthase subur | ATP5O | VVRPFAKLVRPPV | >tr\|H7C086\|H7C086_ | 1865 | | 1 | 3,0138 | 2,1653 | 1,37 | 1,9299 | A |
| B0YJC4;P08670;A0A | 61;61;61;61 | B0YJC4 | B0YJC4 | Vimentin | VIM | PSTSRSLYASSPGG | >tr\|B0YJC4\|B0YJC4_H | 1244 | 1;2 | | 2,0984 | 3,8164 | 1,46 | 1,8265 | A |
| Q9UHR4 | 380 | Q9UHR4 | Q9UHR4 | Brain-specific angio | BAIAP2L1 | DVITLLIPEEKDGW | >sp\|Q9UHR4\|BI2L1_H | 3601 | | 1 | 3,8384 | 2,3725 | 1,5 | 1,9886 | A |

[illegible]

| | |
|---|---|
| **common to the 4 conditions** | **135** |
| **common to 3 conditions** | **64** |
| **common to Src3A** | **6** |
| **common to Src WT** | **3** |
| **unique** | **27** |
| **common to Replicate1** | **19** |
| **common to Replicate2** | **18** |
| **excluded (all conditions <2)** | **7** |
| | **279** |
